# Novel glycoside hydrolase family enzymes from *Escherichia coli* are associated with osmo-regulated periplasmic glucan synthesis

**DOI:** 10.1101/2023.03.29.533101

**Authors:** Sei Motouchi, Kaito Kobayashi, Hiroyuki Nakai, Masahiro Nakajima

## Abstract

Most Gram-negative bacteria synthesize osmo-regulated periplasmic glucans (OPG) in the periplasm or extracellular space. Many pathogens lose their pathogenicity by knocking out *opgG*, an OPG-related gene indispensable for OPG synthesis. However, the biochemical functions of OpgG and OpgD, a paralog of OpgG, have not been elucidated. In this report, structural and functional analyses of OpgG and OpgD from *Escherichia coli* revealed that these proteins are β-1,2-glucanases with remarkably different activity, establishing a new glycoside hydrolase family. Furthermore, a reaction mechanism with an unprecedentedly long proton transfer pathway is proposed for OpgD. The conformation of the region that forms the reaction pathway differs noticeably between OpgG and OpgD, which explains the observed low activity of OpgG. The findings enhance our understanding of OPG biosynthesis and provide insights into functional diversity for this novel enzyme family.

---

Glycans play essential roles as energy sources and in forming structural components such as cell walls and exoskeletons. Glycans also facilitate interactions between organisms to enable processes such as pathogenicity, symbiosis, cell adhesion and signaling. Osmo-regulated periplasmic glucans (OPG), whose carbohydrate moieties are composed of glucose, are synthesized in the periplasm or extracellular region by various Gram-negative bacteria. Although OPG were initially found as glycans synthesized at low osmolarity^1^, various physiological roles of OPG, including pathogenic and symbiotic functions, have been reported. For example, *Xanthomonas campestris*, *Agrobacterium tumefaciens* and *Salmonella enterica* sv. *typhimurium*, regardless of phytopathogens or animal pathogens, lose their pathogenicity by knocking out OPG-associated genes^1, 2^. Some species in Rhizobiaceae use OPG as a symbiotic factor^1, 2^. Thus, OPG are important glycans, especially for enabling interactions between organisms.

OPG are classified into four groups based on their glucan backbone structures. Group 1 has a linear β-1,2-glucosyl backbone with β-1,6-glycosyl branches, whereas groups 2–4 have cyclic backbones (cyclic β-1,2-glucan without side chains, cyclic β-1,2-glucan with one linkage substituted with α-1,6-linkage, and cyclic β-1,3-β-1,6-glucan, respectively)^1^.

Groups 2 and 3 OPG are essentially β-1,2-glucans. Group 2 OPG are found in various species such as *Rhizobiaceae* and *Brucella* and are synthesized by cyclic glucan synthases (Cgs)^3–6^ and transported to the periplasm by Cgt, an ABC transporter^7^. Group 3 OPG are found in *Ralstonia solanacearum*, *Xanthomonas campestris* and *Rhodobacter sphaeroides*^8–10^. Because *R. sphaeroides* produces OPG even if the cgs gene is deleted, genes involved in synthesizing the β-1,2-glucan backbone are unclear^10^. In addition, the enzymatic mechanism that completes cyclization via an α-1,6-glucosidic linkage remains unknown. In contrast to these two groups, Group 4 OPG are formed with β-1,3- and β-1,6-glucosidic linkages by NdvB, which is composed of a glycosyltransferase family 2 domain and a glycoside hydrolase (GH) family 17 domain, and NdvC, which is hypothesized to be a β-1,6-glucan synthase^11–14^.

Group 1 OPG with linear backbones are found in various gamma proteobacteria such as *Escherichia coli*, *Pseudomonas aeruginosa* (an opportunistic pathogen), *Dickeya dadantii* and *Pseudomonas syringae* (phytopathogens)^1^. Hereafter, the word OPG refers to Group 1 OPG. Genetic analysis indicates that the *opgGH* operon is responsible for OPG synthesis. This operon is distributed widely among almost all gamma proteobacteria^15^. Various pathogens lose their pathogenicity by knocking out *opgH* and/or *opgG* genes^1, 2^. Recently, the OpgGH operon in *E. coli* was reported to be related to antibiotic resistance in silkworm^16^. In *D. dadantii* and *S. enterica* sv. *typhimurium*, OPG are known to regulate Rcs phosphorelay directly^2^. Rcs phosphorelay is a key system that regulates the expression of a set of genes encoding plant cell wall-degrading enzymes and the flhDC master operon, the flagellum structural gene of *D. dadantii*^2^.

Chemical structures of OPG have been studied, especially in *E. coli*. Degrees of polymerization (DPs) of OPG are limited to 5–12^1^. OPG can be further modified with phosphoglycerol, succinate and/or phosphoethanolamine groups by OpgB, OpgC and/or OpgE, respectively^17–22^. In contrast to the physiological aspects and chemical structures of OPG, enzymes associated with synthesizing the glucosyl backbone of OPG have not been investigated fully. OpgH was suggested to synthesize a β-1,2-glucosyl main chain enzymatically^23^. However, these analyses were performed using a crude membrane fraction containing recombinant OpgH, and the reaction products were not confirmed biochemically to be β-1,2-glucans. OpgG is hypothesized to play an important role in OPG backbone synthesis because an *opgG* knockout mutant cannot synthesize OPG, regardless of side chains^15^. Although it is postulated that OpgG is involved in forming β-1,6-glucosyl side chains, the ligand-free structure of OpgG from *E. coli* (EcOpgG) provided limited information on detailed biochemical functions^24^. OpgD from *E. coli* (EcOpgD), a paralog of EcOpgG with 32.9% amino acid sequence identity, is another key protein for linear OPG synthesis. Long linear β-1,2-glucosyl main chains with β-1,6-glucosyl side chains were detected in an *opgD* knockout mutant, suggesting that OpgD is related to adjusting chain lengths^25^. Furthermore, the *ΔopgD* mutant loses its motility and flagella, which are restored by deleting Rcs system-related genes^26^. However, no biochemical analysis of OpgD has been performed. Such a lack of biochemical evidence limits our understanding of the mechanism of OPG biosynthesis.

In this report, structural and functional analyses of EcOpgD and EcOpgG were performed. We discovered that EcOpgD is a β-1,2-glucanase (SGL) with a unique reaction mechanism, thereby establishing a new GH family (GHxxx) that is a phylogenetically new group, indicating that novel glycoside hydrolases are involved in OPG biosynthesis. Comparing the EcOpgD structure with EcOpgG with low SGL activity provided insights into the diversity of the functionally important region for hydrolysis by this GH family.

## Results

### Enzymatic properties of EcOpgD and EcOpgG

Recombinant EcOpgD and EcOpgG fused with a His_6_-tag at the C-terminus were produced using *E. coli* as a host and purified successfully. EcOpgD exhibited hydrolytic activity toward β-1,2-glucans to produce Sop_6–7_ preferentially, and Sop_8–10_ accumulated during the late stage of the reaction (Fig. 1a left). Thus, β-1,2-glucans were used as substrates to investigate pH and temperature profiles. EcOpgD was stable at pH 4.0–10.0 and up to 50 °C. EcOpgD exhibited the highest activity at 40 °C and pH 5.0 (Extended Data Fig. 1a–d). These results imply that EcOpgG, a paralog of EcOpgD, also has SGL activity. EcOpgG was found to hydrolyze β-1,2-glucans; however, the catalytic velocity was very low (Fig. 1a *right*). Sop_6–8_ were preferentially produced by EcOpgG, and Sop_9–11_ also accumulated as final products. EcOpgG exhibited the highest activity at pH 5–7 (Extended Data Fig. 1e).

**Fig. 1.**
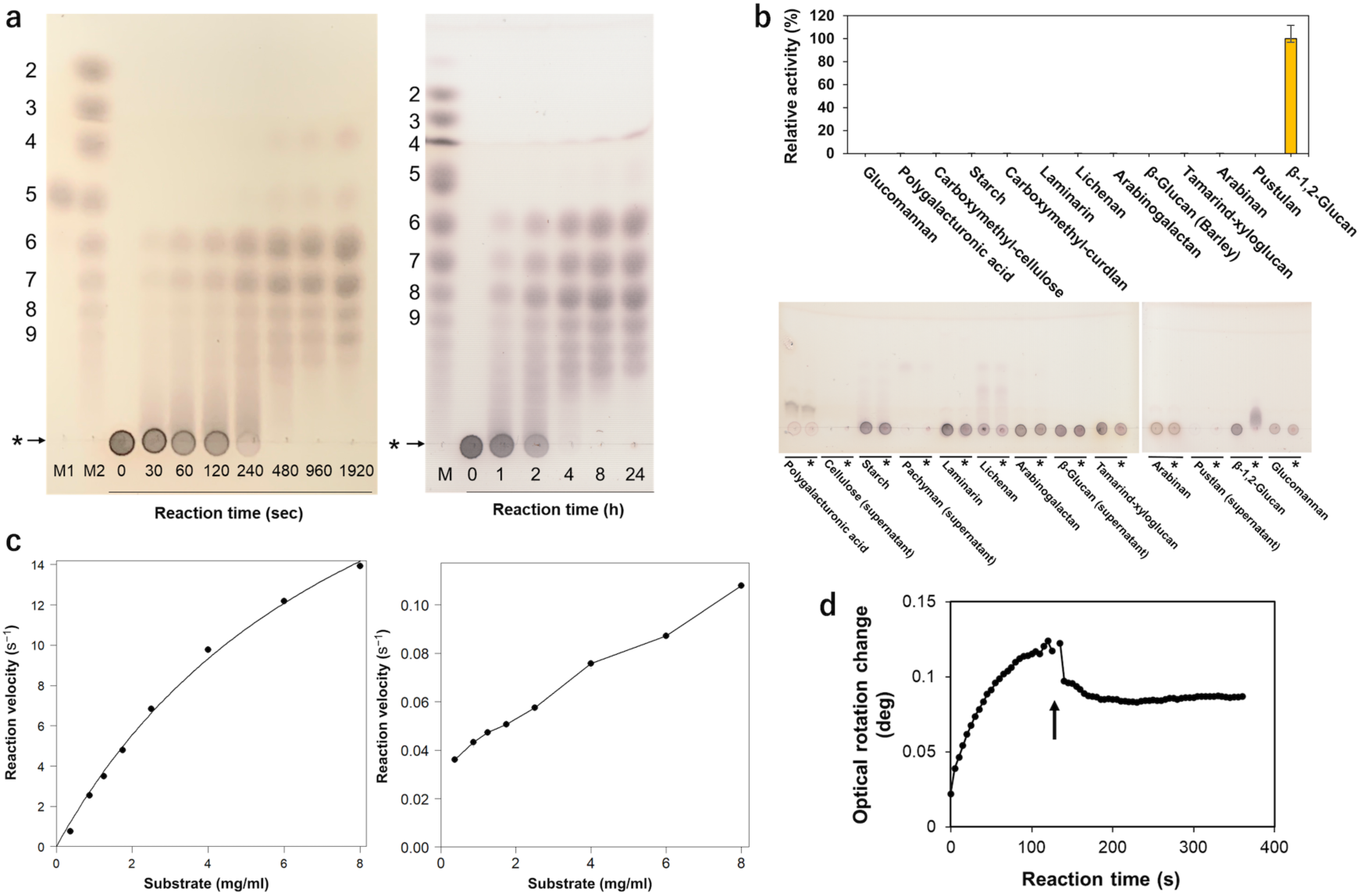
Catalytic analyses of EcOpgD and EcOpgG. **a**, TLC analysis of the action patterns of EcOpgD (*left*) and EcOpgG (*right*) on β-1,2-glucans. Lane M1, 10 mM Sop_5_; Lane M2, a Sop_n_s marker prepared using 1,2-β-oligoglucan phosphorylase from *Listeria inocua*. DPs of Sop_n_s are shown on the left side of the TLC plates. Arrows represent β-1,2-glucan used for reactions. The origins of the TLC plates are shown as horizontal arrows denoted by asterisks. EcOpgD and EcOpgG in the reactions are 0.14 mg/mL and 0.77 mg/mL, respectively. **b**, Substrate specificity of EcOpgD (*top*) and EcOpgG (*bottom*). (*top*) N.D. represents values with less than 0.02% relative activity. (*bottom*) The reaction was performed at 37 °C for 24 h. Asterisks indicate that the reaction time was 24 h. Other lanes represent a reaction time of 0 h. **c**, Kinetic analysis of EcOpgD (*left*) and EcOpgG (*right*). (*left*) Data plotted as closed circles were regressed with the Michaelis–Menten equation (solid line). (*right*) Plots were simply connected by lines because of poor fitting to the Michaelis–Menten equation. **d**, Time course of the observed optical rotation during β-1,2-glucan-hydrolysis by EcOpgD. The arrow indicates that several drops of aqueous ammonia were added to the reaction mixture 120 s after the reaction started.

Among the various tested polysaccharides, EcOpgD exhibited high hydrolytic activity toward β-1,2-glucan but not other substrates, indicating that EcOpgD is highly specific to β-1,2-glucan (Fig. 1b *top*). Thus, kinetic analysis of EcOpgD using β-1,2-glucan as the substrate was performed (Fig. 1c *left*). The *k*_cat_ of EcOpgD was comparable to those of SGLs from *Chitinophaga pinensis* (CpSGL) and *Talaromyces funiculosus* (TfSGL), the first SGLs found in a bacterium and a fungus, respectively (Table 1)^27, 28^. The *K*_m_ of EcOpgD was much higher than those of the two aforementioned SGLs, leading to a much lower *k*_cat_/*K*_m_ for EcOpgD. Nonetheless, the kinetic parameters of EcOpgD are still within the range of those of general GH enzymes, such as the CMCase from *Arcticibacterium luteifluviistationis*^29^.

**Table 1.** Kinetic parameters of EcOpgD for β-1,2-glucans and comparison with known SGLs

| Enzyme | $K_m$ (mg/mL) | $k_{cat}$ ($s^{-1}$) | $k_{cat}/K_m$ ($s^{-1} \text{ mg}^{-1}\text{mL}$) |
| --- | --- | --- | --- |
| EcOpgD | $8.6 \pm 1.1$<br>($0.44 \pm 0.056$)<sup>b</sup> | $29 \pm 2$ | $3.4 \pm 0.2$ |
| CpSGL<sup>a</sup> (GH144) | $0.71 \pm 0.15$<br>($0.068 \pm 0.014$)<sup>b</sup> | $49 \pm 4$ | $69 \pm 10$ |
| TfSGL<sup>a</sup> (GH162) | $0.18 \pm 0.02$<br>($0.015 \pm 0.001$)<sup>b</sup> | $31 \pm 1$ | $170 \pm 10$ |
| EcOpgG | | ($0.11$)<sup>c</sup> | |

<sup>a</sup> The values of CpSGL and TfSGL are cited from Abe et al.<sup>27</sup> and Tanaka et al.<sup>28</sup>, respectively.
<sup>b</sup> The values calculated using molar concentrations are shown in parentheses.
<sup>c</sup> Reaction velocity in the presence of 0.8% β-1,2-glucan at pH 5.5 is shown in parenthesis.

EcOpgG also showed the same substrate specificity toward the polysaccharides as that of EcOpgD. However, the reaction velocity of EcOpgG was low throughout the substrate concentration range examined (Fig. 1b *bottom*, c *left*). The relative activity against EcOpgD was only 0.79% at 8.0 mg/mL β-1,2-glucan (Table 1).

The change in the degree of optical rotation during the hydrolysis of β-1,2-glucans was examined to determine the reaction mechanism of EcOpgD. An increase in the degree of optical rotation during the early stage of the reaction and a rapid decrease in the value upon adding aqueous ammonia were observed (Fig. 1d). This result is similar to observations made for CpSGL and TfSGL, which are inverting SGLs^27, 28^. Thus, EcOpgD follows the anomer inverting mechanism.

### Overall structure of ligand-free EcOpgD

The ligand-free structure of EcOpgD was determined at 2.95 Å resolution (Fig. 2a *left*). A dimer was used for describing the EcOpgD structure because the stable assembly in the structure was found to be a dimer (Fig. 2a *right*). The overall structure of the ligand-free EcOpgD resembles that of the ligand-free EcOpgG.

**Fig. 2.**
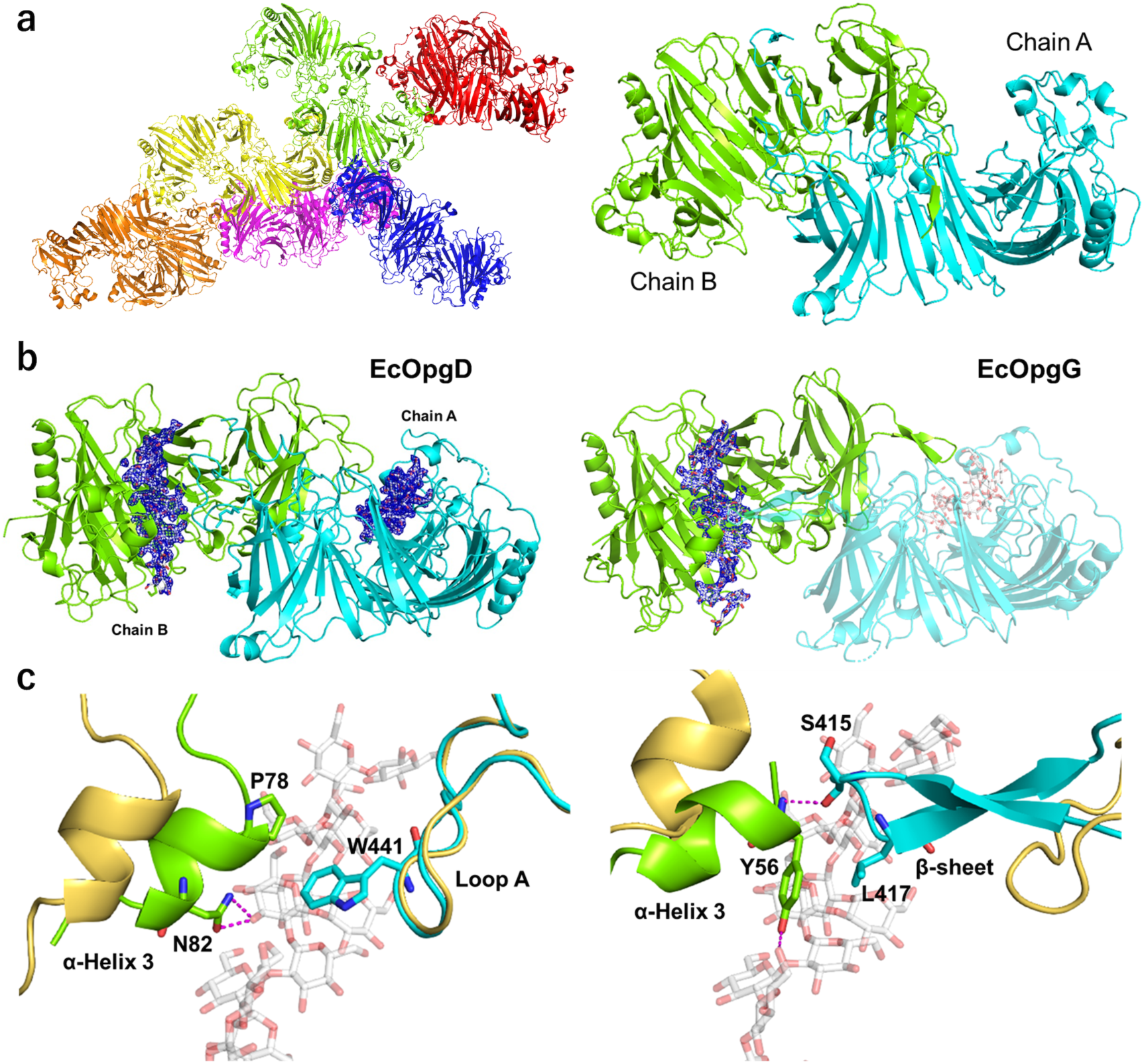
Structures of EcOpgD and EcOpgG. **a,** Overall structure of ligand-free EcOpgD. (*left*) Structure of an asymmetric unit. There are 12 monomers in an asymmetric unit with almost identical conformations (RMSD within 0.30 Å). The free energy of assembly dissociation (Δ*G*_diss_) was calculated by PISA^57^. All Δ*G*_diss_ values of dimer interfaces in the asymmetric unit are higher than 32.9 kcal/mol, indicating that the dimer is a stable assembly. Thus, the six dimers are shown in red, blue, magenta, light green, yellow and orange. (*right*) Biological assembly. Chains A and B are shown in cyan and light green, respectively. The RMSD between EcOpgD and EcOpgG (PDB: 1txk) is 2.206 Å. **b**, Overall structures of Michaelis complexes of the EcOpgD D388N (*left*) and EcOpgG D361N (*right*) mutants. Chains A and B are shown in cyan and light green, respectively. The two subunits represent a biological assembly. The electron densities of β-1,2-glucans are shown as *F*_o_–*F*_c_ omit maps by blue meshes at the 3σ contour level. Substrates are shown as white sticks. (*left*) The biological assembly is identical with molecules in an asymmetric unit (RSMD: 0.170 Å). (*right*) A plausible bioassembly of EcOpgG in solution is a dimer, according to PISA analysis^57^. Chains A and B are a symmetry mate. **c**, Superposition between the ligand-free and the Michaelis complex structure of EcOpgD (*left*) and EcOpgG (*right*) around the Loop A region. The ligand-free structures are shown in light yellow. (*left*) P78, N82 and W441 in the complex structure are shown as sticks. (*right*) Y56, S415 and L417 in the complex structure are shown as sticks.

### Michaelis complex of EcOpgD and EcOpgG

The D388N mutant was used to determine a Michaelis complex structure of EcOpgD because this mutant displayed the lowest activity among mutants of acidic amino acids conserved in its homologs, and this residue is located in the long cleft of the ligand-free structure (Table 2, Extended Data Fig. 2). The complex structure was obtained as a dimer by co-crystallizing the D388N mutant with β-1,2-glucan (Fig.2b *left*). In the catalytic pockets of chains A and B, the electron densities of β-1,2-glucans with DP11 and DP13 (Sop_11_ and Sop_13_, respectively) were clearly observed, respectively (Fig. 2b *left*, Extended Data Fig. 3A). Chain B was used for describing the complex because the glucan chain is longer.

**Table 2.** Specific activities of EcOpgD mutants

|  | Specific activity (U/mg) | Relative activity (%) |
| --- | --- | --- |
| E209Q | 0.48 (0.028) | 3.5 |
| D300N | 0.13 (0.0048) | 0.90 |
| D351N | 3.3 (0.31) | 23 |
| E385Q | 0.28 (0.014) | 2.0 |
| D388N | 0.022 (0.0009) | 0.16 |
| E439Q | 0.77 (0.018) | 5.6 |
| E445Q | 1.9 (0.073) | 14 |
| R359A | 0.064 (0.0013) | 0.46 |
| T386L | 0.78 (0.024) | 5.6 |
| T386A | 0.80 (0.013) | 5.8 |
| W441L | 0.87 (0.013) | 6.3 |
| W441F | 1.0 (0.013) | 7.3 |
| P78L | 8.2 (0.50) | 59 |
| P78A | 6.5 (0.064) | 47 |
| N82A | 12 (0.20) | 87 |
| Y356F | 2.2 (0.054) | 16 |
| Wild-type | 14 (0.33) | 100 |

Medians of triplicate experiments are presented.
Maximum differences from medians are shown in parentheses.

The substrate binding site is formed as a cleft in the ligand-free structure, whereas this site forms a tunnel in the structure of the complex (Fig. 2a, b, Extended Data Fig. 4). The difference arises mainly from the closure motion of α-helix 3 to the substrate in the cleft. In the helix of the complex structure, P68 is located in close proximity to W441, and N82 forms hydrogen bonds with the 3-hydroxy group of the Glc moiety (Fig. 2c *left*). However, substituting P78 (P78A and P78L) and N82 (N82A) only had a mild effect on activity (Table 2), implying that the closure motion of α-helix 3 is not important for catalysis. Unlike the motion of α-helix 3, chain A is involved in the reaction mechanism. In particular, residues 434–453, hereafter termed Loop A, are important for catalysis, as described below. The complex structure of EcOpgG (D361N mutant) with β-1,2-glucan was determined at 1.81 Å resolution to delineate the very low reaction velocity of EcOpgG (Fig. 2b *right*). In the catalytic pocket of the complex, the electron density of a Sop_16_ molecule was observed (Extended Data Fig. 3b). The dynamic conformational change from a loop (residues 409–425) to two β-strands upon substrate binding is a unique feature found in EcOpgG (Fig. 2c *left*). However, the β-sheet interacts not with the substrate but with the α-helix 3 in the complex. The motion of α-helix 3 is induced by the interactions between Y56 in the helix and the 6-hydroxy group of the Glc moiety. These observations suggest that the closure motion of α-helix 3 is required for forming the β-sheet. In contrast, Loop A in EcOpgD does not change its conformation upon substrate binding nor does it have electrostatic interactions with α-helix 3 (Fig. 2c *left*). Because α-helix 3 in EcOpgD is not required for catalytic activity (see above), the motion of this helix may be a remnant of molecular evolution from EcOpgG.

### Substrate binding mode of EcOpgD and EcOpgG

We searched for distorted Glc moieties to determine the position of the glycosidic bond cleaved. Subsite –1 is often distorted to enable a nucleophile to attack an anomeric center. The seventh and eleventh Glc moieties from the reducing end of Sop_13_ form skew boat (^1^S_3_ and ^1^S_5_, respectively) conformations, according to Cremer–Pople parameters^30^. The other Glc moieties form a chair (^4^C_1_) conformation. A water molecule is located near the anomeric carbon of the seventh Glc moiety from the reducing end (3.5 Å). The angle formed by this water, the anomeric carbon and the glycosidic bond oxygen atom is 168.17°, which is close to 180° and suitable for the nucleophilic (in-line) attack of the anomeric carbon (Fig. 3a). The position of the water molecule is consistent with the result that EcOpgD follows an anomer inverting mechanism. The other Glc moiety with the skew boat conformation has no nucleophilic water because the O5 atom of the twelfth Glc moiety from the reducing end occupies the position for a nucleophile. These observations strongly suggest that the position of the seventh Glc moiety is subsite –1 and that EcOpgD accommodates a Sop_13_ molecule at subsites –7 to +6 as a Michaelis complex (Extended Data Fig. 5a). This is consistent with the preferential production of Sop_6–7_ by hydrolysis of β-1,2-glucans (Fig. 1a *left*).

**Fig. 3.**
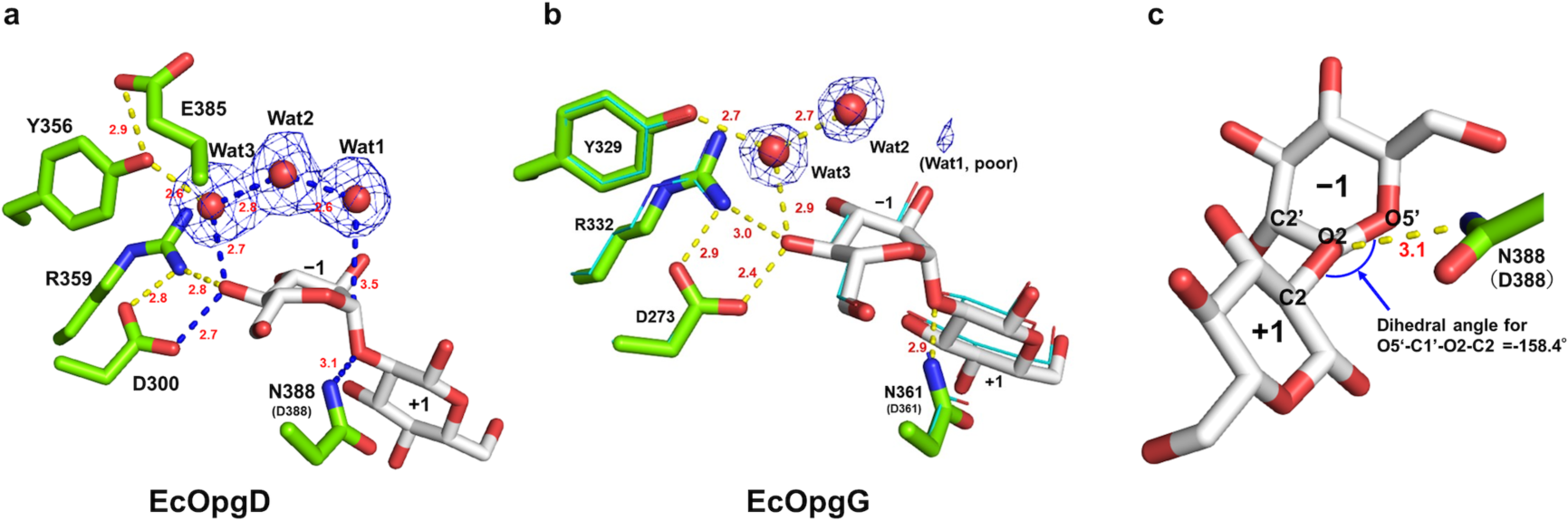
Substrate binding modes of EcOpgD and EcOpgG complex structures. Hydrogen bonds with distances (Å, red numbers) are shown as yellow dotted lines. **a**, **b**, Structures around subsite –1 in EcOpgD (**a**) and EcOpgG (**b**). Hydrogen bonds on the suggested reaction pathway are shown as blue dotted lines. The electron densities of Wat1, Wat2 and Wat3 are shown as *F*_o_–*F*_c_ omit maps by blue meshes at the 3σ contour level. Residues and substrates are shown in green and white sticks, respectively. **b**, EcOpgD (cyan, line representation) superimposed with EcOpgG. **c**, Conformation of the Glc moieties at subsites –1 and +1 in EcOpgD. The innate wild-type residues are shown in parentheses if the residues are substituted.

At least two residues participate in the interaction with Glc moieties at subsites –5 to +4. In particular, the Glc moiety at subsite –1 is recognized by five residues, which allows the distorted conformation. The Glc moieties at subsites –6, +5 and +6 are mainly recognized intra-molecularly rather than forming hydrogen bonds with nearby residues (Supplemental Table S1, Extended Data Fig. 5a).

A β-1,2-glucan molecule binds to the substrate pocket in the complex of EcOpgG at almost the same position as EcOpgD, except at the subsite minus side from –5 in EcOpgD. Substrate recognition residues are well conserved at subsites –1 to +4 between the two enzymes but are not at other subsites, especially on the minus side (Supplemental Table S1, Extended Data Fig. 5b). Although the conformation of the Glc moiety at subsite –5 in EcOpgD is ^1^S_5_, the corresponding Glc moiety in EcOpgG forms the ^4^C_1_ conformation. The only Glc moiety forming ^1^S_3_ in EcOpgG is the well-superimposed Glc moiety at subsite –1 in EcOpgD. However, electron density for nucleophilic water is poor in EcOpgG, which strongly supports the observed low SGL activity of EcOpgG (Fig. 3b).

### Catalytic residues of EcOpgD

Generally, GH enzymes have two acidic residues as catalysts. D388 was identified as a general acid catalyst candidate because this acidic amino acid directly interacts with the oxygen atom of the glycosidic bond between subsites –1 and +1 (Fig. 3a). The distance between the nitrogen atom of the N388 side chain and O2 atom is 3.1 Å, and the dihedral angle for O5’–C1’–O2–C2 is within the range of antiperiplanar (–158.4°) (Fig. 3c), suggesting that the substrate and D388 are properly arranged for *syn* protonation^31, 32^. The relative activity of D388N was the lowest in all examined mutants (0.15%) (Table 2). These results strongly suggest that D388 is a general acid residue.

No acidic residue directly interacting with the nucleophilic water is found in the complex structure, unlike canonical anomer inverting GH enzymes. Thus, proton dissociable amino acid residues (D300, Y356, R359 and E385) on possible proton transfer pathways from the nucleophilic water were listed as general acid candidates (Fig. 3a). All listed residues are via at least three water molecules, including the nucleophilic water. Hereafter, these water molecules are called Wat1, Wat2 and Wat3 from the nucleophilic water. Although Wat2 interacts with no proton dissociative residues, Wat3 interacts directly with Y356. The relative activity of Y356F was 14% (Table 2), suggesting that Y356 is not a general base. This result also suggests that E385 beyond Y356 is not a general base despite the dramatic decrease in activity observed for the E385Q mutant (Fig. 3a, Table 2). D300 and R359 interact with Wat3 via the 4-hydroxy group in the Glc moiety at subsite –1. The relative activities of D300N and R359A were 0.89% and 0.47%, respectively (Table 2). The predicted p*K*_a_ values by propKa 3.4.0^33^ for the carboxy group of D300 and guanidium group of R359 are 4.1 and 13.8, respectively. An electrostatic interaction between D300 and R359 probably promotes deprotonation of the D300 carboxy group as a general base and protonation of the guanidium group of R359. The p*K*_a_ value of the D388 carboxy group substituted with asparagine in the complex structure of the D388N mutant was predicted to be 5.5, which is consistent with the optimum pH 5.0 between the predicted p*K*_a_ values of the two carboxy groups in D300 and D388. Overall, a nucleophilic water is likely activated by D300 as a general base through the two water molecules (Wat2 and Wat3) and one substrate hydroxy group.

### Environment around the water molecules on the proposed catalytic pathway

The high efficiency of proton relays that carry a proton from a nucleophilic water to a general base via other water molecules has been explained using the Grotthuss mechanism in GH enzymes. To further judge the plausibility of the suggested proton transfer pathway, the environment around Wat1 (nucleophilic water), Wat2 and Wat3 was examined, and mutational analysis was performed. The three water molecules are sequestered from the solvent, especially by the β-1,2-glucan and Loop A. Other loops (residues 349–355 and 380–389 of chain B) also participate in sequestering these water molecules from the solvent.

The nucleophilic water is surrounded by W441 and T386 (Fig. 4a, b). Substitution of W441 with Phe or Leu, smaller hydrophobic side chain amino acids, resulted in greatly reduced relative activities (7.7% and 6.5%, respectively) (Table 2). Interaction of the hydroxy group of T386 with E445 causes the methyl group of T386 to face the nucleophilic water. The relative activity of T386S, a mutant without a methyl group, decreased (17.5% relative activity) (Table 2). These results suggest that the hydrophobicity provided by the six-membered ring moiety of W441 and the methyl group of T386 around the nucleophilic water support efficient proton transfer by the Grotthuss mechanism. Wat2 is surrounded by the Cβ of E385, C5 of the Glc moiety at subsite –1 and the main chain oxygen atom of G440 (Fig. 4a, b). The G440 oxygen atom forms a hydrogen bond with Wat2, suggesting that G440 fixes Wat2 to an appropriate position for intermediate proton transfer from Wat1 to Wat3. Wat3 is sequestered from the solvent by the 6-hydroxy group of the Glc moiety at subsite –3, the 4-hydroxy group of the Glc moiety at subsite –1, Y356 and R359 (Fig. 4a, c). Wat3 forms hydrogen bonds with the 6-hydroxy group of the Glc moiety at subsite –3 and the hydroxy group of Y356. The reduced activity of Y356F (16% relative activity) suggests that Y356 is not a general base, as described above, but its hydroxy group is important for fixing and orienting Wat3 (Table 2).

**Fig. 4.**
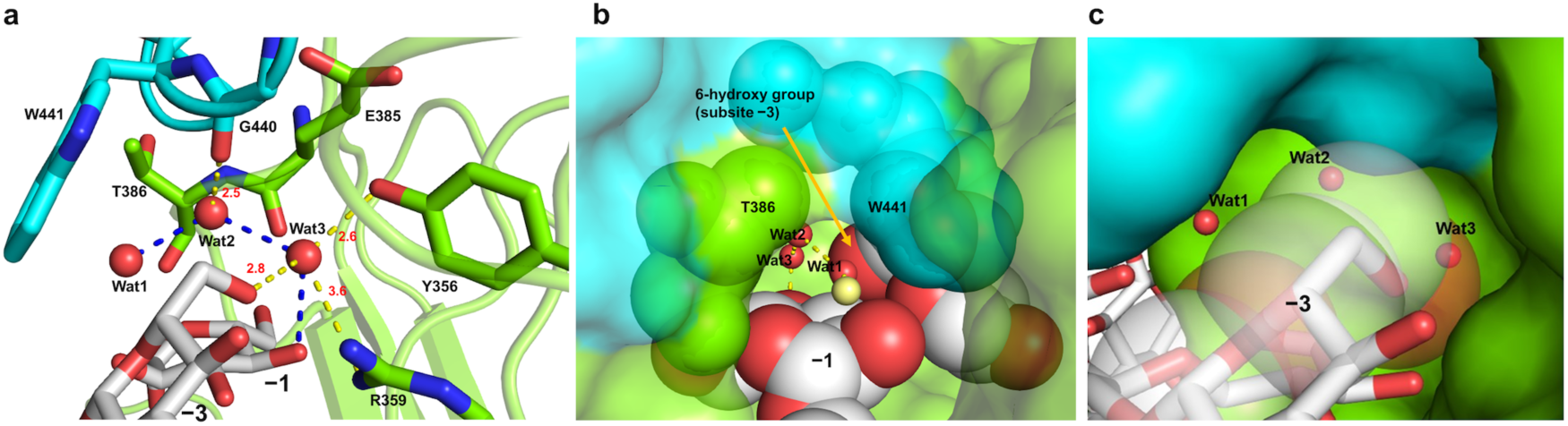
Water molecules are indispensable for the reaction pathway. Chains A and B are shown in cyan and light green, respectively. Hydrogen bonds with distances (Å, red numbers) are shown as yellow dotted lines. Substrates are shown as white sticks. **a**, Residues sequestering water molecules on the reaction pathway. All residues used for fixing Wat1–3 on the reaction pathway are shown as sticks. Blue dotted lines represent a route from the nucleophile to a substrate hydroxy group on the reaction pathway. **b**, **c**, The environment around the sequestered Wat1 (**b**) and Wat2 and Wat3 (**c**). **b**, The substrate and T386 and W441 are shown as spheres with van der Waals radii. **c**, The substrate is shown in stick representation. C1, C5, O5 and O6 atoms of the Glc moiety at subsite –3 are shown as semi-translucent spheres with van der Waals radii.

Loop A, important for sequestering water molecules, is tethered by interactions involving E385, E439 and E445 (Extended Data Fig. 6). The relative activities of the E385Q, E439Q and E445Q mutants were greatly reduced (2.0%, 5.6%, 14%, respectively) (Table 2). Even substituting glutamate residues with similar amino acids affected hydrolytic activity noticeably, indicating the vital role of Loop A in the catalytic mechanism of EcOpgD. Overall, structural data and mutational analyses support the suggested proton transfer pathway. We summarize the proposed reaction mechanism of EcOpgD in Fig. 5.

**Fig. 5.**
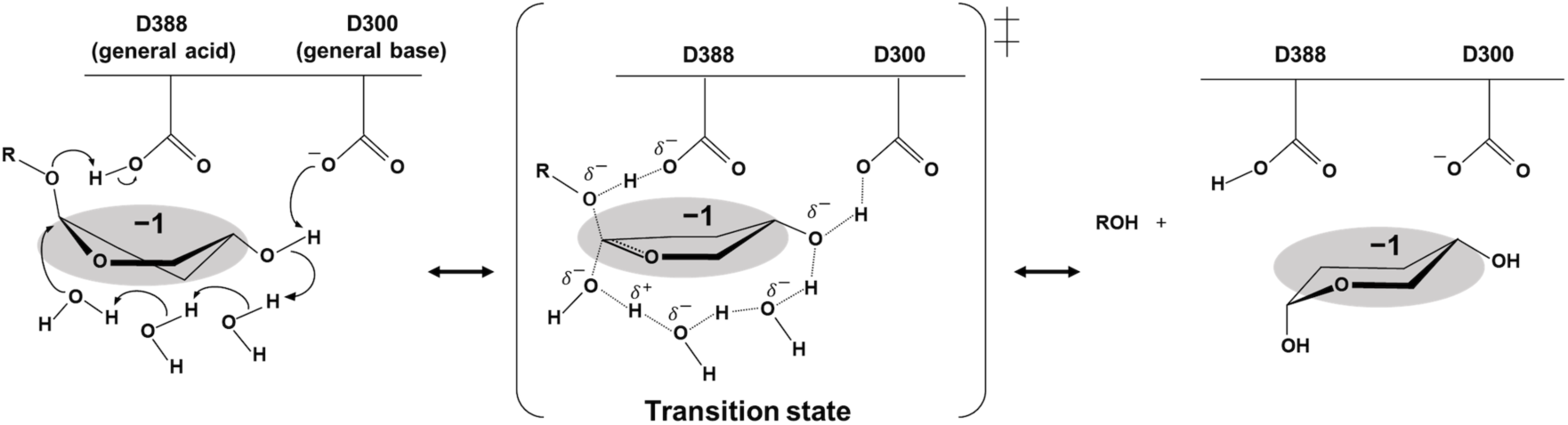
A proposed reaction mechanism of EcOpgD. The Glc moiety at subsite –1 is highlighted in gray. Arrows represent the pathway for electron transfer.

### Comparison of the catalytic mechanism with EcOpgG

Intriguingly, residues associated with catalysis in EcOpgD (D388, R359, D300) are conserved in EcOpgG (Fig. 3a, b, Extended Data Figs. 2, 5c), although the reaction velocity of EcOpgG was very low (Fig. 1c *right*). This result indicates significant differences in the environments around the reaction route between EcOpgG and EcOpgD. In fact, a poor nucleophilic water was observed in EcOpgG, as described above (Fig. 3b). This difference in activity appears to be caused by the hydrophobic amino acids surrounding the nucleophilic water. W441 in EcOpgD is spatially substituted with L417 in EcOpgG (Figs. 2c, 6), whereas T386 in EcOpgD is conserved in EcOpgG (Extended Data Figs. 2, 5c). The other difference is the environment around Wat2. In EcOpgG, two water molecules form a route to facilitate the escape of Wat2 to solvent, whereas G440 in EcOpgD blocks this route completely (Fig. 6). These observations explain the low activity of EcOpgG toward β-1,2-glucans and strongly support the suggested proton transfer pathway of EcOpgD.

**Fig. 6.**
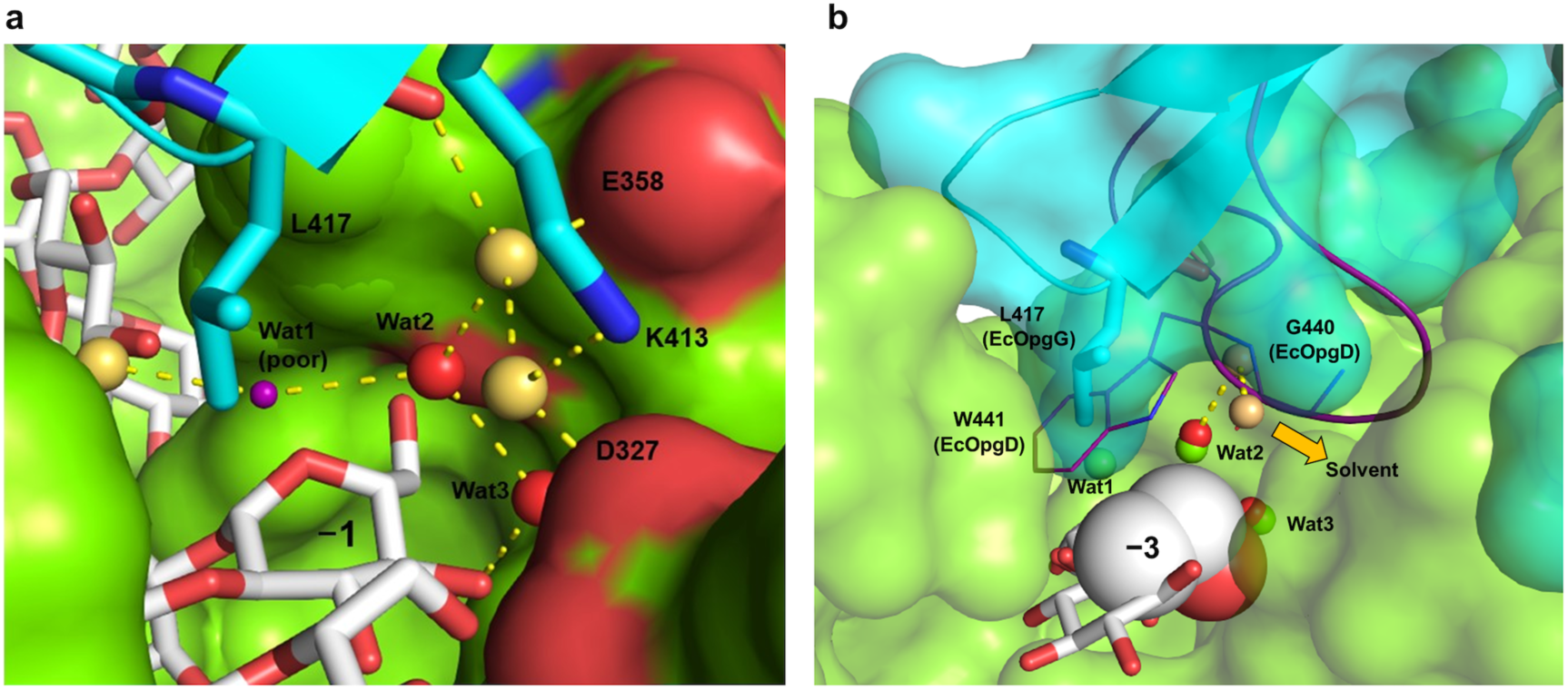
The environment around Wat2 and Wat3 in the EcOpgG complex structure. **a**, The positions of the water molecules around subsite –1 of EcOpgG. A small purple sphere is shown as a tentative water molecule with poor electron density (Wat1). Wat2 and Wat3 and the other water molecules are shown in red and beige spheres, respectively. K413 and L417 are shown as cyan sticks. Chains A and B are shown in blue cartoon and green surface representations, respectively. The substrate is shown as a white stick. **b**, Superposition showing the spatial position of Loop A in EcOpgG and EcOpgD. The superimposed Loop A in the EcOpgD complex is shown in purple. G440 and W441 are shown as lines. Wat1–3 molecules in EcOpgD are shown as light green spheres. EcOpgG is shown as a semi-translucent surface. L417 is shown in cyan. The water molecules of EcOpgG are shown as presented in (**a**), except that Wat1 (poor) is omitted. The Glc moieties at subsites –1 and –3 of EcOpgG are shown as white sticks. C5, C6 and O6 atoms of the Glc moiety at subsite –3 in EcOpgG are shown as spheres of van der Waals radii. The thick yellow arrow represents another proton network that leads to the solvent.

## Discussion

EcOpgG and EcOpgD have been classified at the genetic level as proteins indispensable for OPG biosynthesis and regulation of OPG chain lengths, respectively^15, 25^. Although the shape of the ligand-free EcOpgG cleft suggests binding to glycans^24^, the enzymatic functions of EcOpgD and EcOpgG have not been investigated. This is probably because a canonical reaction mechanism to act on β-1,2-glucans cannot be assumed from the ligand-free structure of EcOpgG based on arrangements of two acidic amino acids as catalytic residues for general GH enzymes. In this report, EcOpgD is shown to be an SGL, strongly suggesting that EcOpgD can adjust OPG chain lengths enzymatically. There are sufficient spaces to accommodate β-1,6-glycosyl side chains at subsites +3, +5 and +6, suggesting that EcOpgD can interact with β-1,6-branched β-1,2-glucans as substrates (Extended Data Fig. 7). In addition, EcOpgG has sufficient spaces at the same subsites as EcOpgD, and additionally at subsites –9, –8, –7 and –5 (Extended Data Fig. 7).

EcOpgG displayed much lower SGL activity than EcOpgD. However, the catalytic residues and R359 supporting deprotonation of D300 are highly conserved in OpgG and OpgD homologs. Moreover, the distorted Glc in the Michaelis complex of EcOpgG structurally resembles that of EcOpgD at subsite –1 (Fig. 3b), suggesting that β-1,2-glucan can bind to EcOpgG with a suitable conformation for hydrolysis. However, the absence of a nucleophilic water molecule and the presence of a branched proton transfer pathway from Wat 2 may be responsible for the low specific activity of EcOpgG (0.11 U/mg). Overall, EcOpgG is intrinsically a GH enzyme but may require a cofactor to display greater specific activity. OpgH has often been predicted to be a cofactor of OpgG because *opgG* and *opgH* form a gene cluster^24^. However, *opgI* and *opgH* from *R. sphaeroides* complement *ΔopgH E. coli* even though the sequences of the periplasmic region are diverse among OpgH homologs^34^, which suggests that OpgG has a periplasmic cofactor other than OpgH.

According to the annotation by InterPro^35^, EcOpgG and EcOpgD belong to the MdoG superfamily, which comprises the MdoD and MdoG families (MdoG and MdoD are the former names of OpgG and OpgD, respectively). Although this superfamily can be divided into 14 clades (Extended Data Fig. 8), members of both families are mixed phylogenetically based on the annotation, which generates a degree of confusion. Highly conserved residues among the MdoG superfamily are located in the substrate pocket (Extended Data Fig. 9a), which contains the catalytic residues identified in this study (Extended Data Fig. 9b). Many of the conserved residues contribute strongly to binding of a β-1,2-glucan molecule, according to estimates of the Gibbs free energy for binding (Extended Data Fig. 9c). In contrast, the remarkable difference in biochemical functions between EcOpgG and EcOpgD was attributed to a region of Loop A. In particular, W441 is important for the fixation of nucleophilic water in EcOpgD; however, this amino acid and G440 are conserved in a limited group, clade 2, containing EcOpgD (Extended Data Fig. 9d). In EcOpgG, L417 is spatially equivalent to W441 in EcOpgD and is conserved only in close homologs of EcOpgG (clade 14). Comparing the Loop A region between clades, the GXGG motif shared among clades 1–4 may represent a functionally common structure (Extended Data Fig. 9c). No other consensus region or residue was identified. Furthermore, some clades have no region corresponding to Loop A, indicating diverse biochemical functions for this superfamily. Knockout of OPG-related genes drastically alters phenotypes related to pathogenicity in many species. Homologs from such species in the MdoG superfamily are found only in clades 2–4 and 14, whereas there is no report on the functions of homologs in clades 5–13. Our report on the structure–function relationships of EcOpgD and EcOpgG will provide important clues to investigate relationships between the enzymatic functions of OPG-related genes and the phenotypes of these species.

GHs are classified into families in the CAZy database based on their amino acid sequences^36^ (http://www.cazy.org/). To understand the category of the MdoG superfamily, we initially compared EcOpgD with the GH144 and GH162 family SGLs, which are all known SGLs. EcOpgD showed neither amino acid sequence similarity nor tertiary structure similarity with GH144 and GH162 SGLs^27, 28^. Performing a BLAST search using EcOpgD as a query against the database containing all GH families gave no GH family enzyme hit with sufficient amino acid sequence identity. According to a DALI (http://ekhidna.biocenter.helsinki.fi/dali_server/)^37^ search, GH38 α-mannosidase (PDB: 2wyh) was the top hit among GH family enzymes. However, only the C-terminal domain, but not the catalytic domain of the GH38 enzyme, showed structural similarity with EcOpgD. Therefore, the MdoG superfamily represents a new GH family, GHxxx.

We also proposed a unique reaction mechanism of EcOpgD (Fig. 5). The proton transfer pathway from a nucleophilic water to a general base is unprecedently long, involving two water molecules and a substrate hydroxy group. No pathway to deprotonate nucleophilic water by a general base via a substrate like EcOpgD has been found in GH families. The number of water molecules passing through is also unprecedented. Nevertheless, the reaction model is credible because each feature is found in known reaction mechanisms. In GH130 and GH162, protonation of a scissile bond oxygen atom is carried out by a general acid via a substrate hydroxy group^28, 38, 39^. In GH6, GH101, GH136 and GH162^28, 40–42^, only one water molecule is inserted in the proton transfer pathway from a nucleophilic water to a general base. The findings in this study expand the diversity of GH enzymes in terms of reaction mechanisms and phylogenetic groups.

We identified the enzymatic function of EcOpgD and suggested a unique reaction mechanism of the enzyme. In contrast, EcOpgG was found to show low SGL activity but have a substrate binding mode suitable for hydrolysis. Intriguingly, comparing these two enzymes revealed that amino acid sequences in the Loop A region are not conserved in the MdoG superfamily, suggesting diverse reaction mechanisms for this family. Discovery of novel GH enzymes involved in OPG biosynthesis and elucidation of reaction mechanisms based on three-dimensional structures underpin future efforts that will focus on understanding OPG biosynthesis. This study should lead to unveiling more interactions between organisms through OPG and facilitate efforts to control OPG biosynthesis in Gram-negative bacteria for regulating pathogenicity and symbiosis.

## Methods

### Cloning and purification of EcOpgD and EcOpgG

Genes encoding EcOpgG (GenBank: C569.1) and EcOpgD (GenBank: C910.2) were amplified by PCR with primer pairs shown in Supplemental Table 3 using PrimeSTAR Max and a colony of *E. coli* BL21(DE3) as the template. The forward primers were designed to eliminate the signal sequence predicted by the signalP5.0 server^43^. The amplified genes were inserted between the XhoI and NdeI sites of the pET30a vector by the SLiCE method^44^ to add a C-terminal His_6_-tag to the target proteins. The constructed plasmids were transformed into *E. coli* BL21(DE3), and the transformants were cultured in 1 L Luria-Bertani medium containing 30 mg/L kanamycin at 37 °C until the absorbance at 600 nm reached 0.6. Expression was then induced by adding isopropyl β-D-1-thiogalactopyranoside to a final concentration of 0.1 mM, and cells were cultured at 20 °C (EcOpgD) or 7 °C (EcOpgG) for a further 24 h. The cells were centrifuged at 7000 *g* for 10 min and suspended in 50 mM Tris-HCl buffer (pH 7.5) containing 50 mM NaCl (buffer C). The suspended cells were disrupted by sonication and centrifuged at 33000 *g* for 15 min to obtain a cell extract. The cell extract was applied onto a HisTrap™ FF crude column (5 ml; GE Healthcare) pre-equilibrated with a buffer containing 50 mM Tris-HCl (pH 7.5), 500 mM NaCl and 20 mM imidazole. After the column had been washed with the same buffer, the target protein was eluted with a linear gradient of 20–300 mM imidazole in a buffer containing 50 mM Tris-HCl (pH 7.5) and 500 mM NaCl. Amicon Ultra 30,000 molecular weight cutoff centrifugal filters (Millipore) were used to concentrate a portion of the fractionated protein and to exchange the buffer to 50 mM Tris-HCl (pH 7.5) containing 50 mM NaCl. Each purified protein migrated as a single band of 60 kDa on SDS-PAGE gels, which is consistent with the theoretical molecular mass of EcOpgD (60601 Da) or EcOpgG (56543 Da).

### β-1,2-Glucans used for experiments

β-1,2-Glucans with an average DP of 121 calculated from their number average molecular weight (Mn) were treated with NaBH_4_ for the MBTH (3-methyl-2-benzothiazolinonehydrazone) assay^45^. The average DP of β-1,2-glucans used for TLC analysis and crystallization to obtain the Michaelis complexes of EcOpgD and EcOpgG were 121, 121 and 17.7 based on Mn, respectively. The average DP of β-1,2-glucans used for polarimetric analysis was estimated to be approximately 25, according to the preparation method^46^.

### MBTH method

Quantification of reaction products released by enzymatic reactions was performed by the MBTH method^47^. After each reaction, 20 μL aliquots of the reaction mixtures were taken and heated at 100 °C for 5 min. The samples were mixed with 20 μL of 0.5 N NaOH and then with 20 μL MBTH solution composed of 3 mg/mL MBTH and 1 mg/mL dithiothreitol. The mixtures were incubated at 80 °C for 30 min. A solution comprising 0.5% FeNH_4_(SO_4_)_2_, 0.5% H_3_NSO_3_ and 0.25 N HCl (40 μL) was added to the mixtures, and then 100 μL distilled water was also added after cooling to room temperature. The absorbance at 620 nm was measured. Sop_2_ (0.3–2.0 mM) was used as the standard.

### TLC analysis

EcOpgD and EcOpgG were incubated with 1% β-1,2-glucan in 25 mM sodium acetate buffer (pH 5.0) and sodium acetate-HCl buffer (pH 5.5, adjusted with HCl), respectively, at 30 °C. After heat treatment at 100 °C for 5 min, the reaction mixtures (1 μL) were spotted onto TLC Silica Gel 60 F_254_ (Merck Millipore) plates. The plates were developed with 75% acetonitrile twice or more. The plates were then soaked in a 5% (w/v) sulfuric acid/methanol solution and heated in an oven until the spots were clearly visualized. A β-1,2-glucooligosaccharides marker was prepared by incubating Sop_3–7_ and β-1,2-glucan in 1 mM sodium phosphate containing 1,2-β-oligoglucan phosphorylase from *Listeria innocua*^48^.

### General properties

To determine the optimum pH, EcOpgD (18.0 μg/ml) was incubated in various 20 mM buffers (sodium acetate-HCl, pH 4.0–5.5; Bis-Tris-HCl, pH 5.5–7.5; Tris-HCl, pH 7.5–9.0; glycine, pH 9.0–10.0) containing 0.8% β-1,2-glucan (treated with NaBH_4_) at 30 °C for 10 min and then heated at 100 °C for 5 min to terminate the reaction. The reducing power of Sop_n_s released from the substrate was measured by the MBTH method^47^. The optimum temperature was determined by performing the reactions in 20 mM sodium acetate buffer (pH 5.0) at each temperature (0–70 °C). The pH stability of EcOpgD was determined by incubating the purified enzyme (0.29 mg/mL) in various 20 mM buffers at 30 °C for an hour, and then the reaction was carried out in 50 mM acetate-Na buffer (pH 5.0) containing 0.8% β-1,2-glucan at 30 °C for 10 min. The temperature stability of EcOpgD (14.4 μg/mL) was determined by incubating the enzyme in 50 mM sodium acetate buffer (pH 5.0) at each temperature (0–70 °C) for an hour, and then the reaction was carried out under the same conditions as used for pH stability testing. The optimum pH of EcOpgG (0.48 mg/mL) was determined by incubating the enzyme with 0.8% β-1,2-glucan (treated with NaBH_4_) in various 50 mM buffers (sodium acetate-HCl, pH 4.0–5.5; MES-NaOH, pH 5.5–6.5; MOPS-NaOH, pH 6.5–7.5; Tris-HCl, pH 7.5–9.0; glycine, pH 9.0–10.0) at 30 °C for 15 min and then heated at 100 °C for 5 min to terminate the reaction. The reducing power of Sop_n_s released from β-1,2-glucans was measured by the MBTH method^47^.

### Substrate specificity

EcOpgD (3.6 μg/mL) was incubated in 50 mM sodium acetate buffer (pH 5.0) containing each substrate (0.05% glucomannan, Megazyme (Wicklow, Ireland); 0.0025% polygalacturonic acid, Megazyme; 0.1% carboxymethyl cellulose, Merck; 0.02% soluble starch, Fujifilm; 0.005% carboxymethyl curdlan, Megazyme; 0.0125% laminarin, Merck; 0.0125% lichenan, Megazyme; 0.006% arabinogalactan, Megazyme; 0.2% barley β-glucan, Megazyme; 0.2% tamarind-xyloglucan, Megazyme; 0.1% arabinan, Megazyme; 0.05% pustulan, InvivoGen; or 0.8% β-1,2-glucan) at 30 °C for 10 min, and then the reducing power of products released from each substrate was measured by the MBTH method. Glc was used as a standard for the assay using substrates, except for β-1,2-glucan. EcOpgG (0.769 mg/mL) was incubated in 50 mM acetate-Na-HCl buffer (pH 5.5) containing 0.25% of each substrate (glucomannan, polygalacturonic acid, cellulose, starch, pachyman, laminarin, lichenan, arabinogalactan, barley β-glucan, tamarind-xyloglucan, arabinan, pustulan or β-1,2-glucan) at 37 °C for 24 h, and reaction patterns were then analyzed by TLC. The supernatants, after flush centrifugation of the reaction solutions for removal of insoluble substrates, were spotted on a TLC plate.

### Kinetic analysis

The kinetic parameters for β-1,2-glucans were determined by performing the enzymatic reaction in a 20 μL reaction mixture containing 18.0 μg/mL EcOpgD, 0.0375–0.8 mM β-1,2-glucan (treated with NaBH_4_) and 20 mM acetate-Na-HCl (pH 5.0) at 30 °C for 10 min. For EcOpgG, 0.48 mg/mL EcOpgG was used for the reaction in 50 mM sodium acetate-HCl (pH 5.5) for 15 min. Color development of the reaction mixtures was performed using the MBTH method^47^. Molar concentrations of β-1,2-glucans were calculated based on Mn. Kinetic parameters of EcOpgD were determined by fitting experimental data to the Michaelis–Menten equation, *v*/[E]_0_ = *k*_cat_ [S]/(*K*_m_+ [S]), where *v* is the initial velocity, [E]_0_ is the enzyme concentration, [S] is the substrate, *k*_cat_ is the turnover number and *K*_m_ is the Michaelis constant.

### Polarimetric analysis of the reaction products of EcOpgD

The time course of the degree of optical rotation in the reaction mixture was monitored to determine the reaction mechanism of EcOpgD. The degree of optical rotation of the reaction mixture containing EcOpgD (0.46 mg/mL) and 2% β-1,2-glucans was measured using a Jasco p1010 polarimeter (Jasco) at room temperature. Several drops of 25% aqueous ammonia were added 120 s after the reaction started to enhance mutarotation between the anomers.

### Crystallography

EcOpgD (D388N) and EcOpgG (D361N) were purified using a HisTrap™ FF crude column, as described above. The ligand-free crystals of EcOpgD for data collection were obtained at 20 °C after 3 days by mixing 1 μL D388N mutant (7.0 mg/mL) and 1 μL reservoir solution containing 0.1 M Tris-HCl buffer (pH 8.5), 0.1 M TMAO and 17% (w/v) PEG 2000 MME. The crystals of EcOpgD for the β-1,2-glucan complex were obtained at 20 °C after several months by mixing 7 mg/mL of the D388N mutant with 1 μL 0.5% β-1,2-glucan and 1 μL reservoir solution comprising 0.1 M MMT (pH 4.0) and 20% (w/v) PEG 1500. The crystal of EcOpgG for the β-1,2-glucan complex was obtained at 20 °C after a day by mixing 1 μL of the D361N mutant (7 mg/mL) and 1 μL reservoir solution containing 0.1 M MMT (pH 5.0) and 27% (w/v) PEG400. Ligand-free crystals of D388N were soaked in the reservoir solution supplemented with 27% (w/v) trehalose. The complex crystals of D388N were soaked in the reservoir solution supplemented with 22% (w/v) PEG 200 and 0.5% (w/v) β-1,2-glucan. The ligand-free crystals of D361N were soaked in a reservoir solution supplemented with 3% (w/v) β-1,2-glucan. Each crystal was kept at 100 K in a nitrogen-gas stream during data collection. All X-ray diffraction data were collected on a beamline (BL-5A) at Photon Factory (Tsukuba, Japan). The diffraction data for the ligand-free EcOpgD crystals, crystals of the β-1,2-glucan-bound EcOpgD and EcOpgG mutants were collected at 1.0 Å and processed with X-ray Detector Software (http://xds.mpimf-heidelberg.mpg.de/)^49^ and the Aimless program (http://www.ccp4.ac.uk/). The initial phases of EcOpgD and EcOpgG structures were determined by molecular replacement using the Alphafold2 predicted EcOpgD and the ligand-free EcOpgG (PDB: 1txk) as model structures, respectively. Molecular replacement, auto model building and refinement were performed using molrep, buccaneer, refmac5 and coot programs, respectively (http://www.ccp4.ac.uk/)^50–53^. Crystallographic data collection and refinement statistics are summarized in Supplemental Table S2. All visual representations of the structures were prepared using PyMOL (https://pymol.org/2/).

### Mutational analysis

The plasmids of EcOpgD and EcOpgG mutants were constructed using a PrimeSTAR mutagenesis basal kit (Takara Bio), according to the manufacturer’s instructions. PCRs were performed using appropriate primer pairs (Supplemental Table S3) and the EcOpgD or EcOpgG plasmid as a template. Transformation into *E. coli* BL21(DE3) and the expression and purification of EcOpgD and EcOpgG mutants were performed using the same method described for wild-type EcOpgD preparation. The enzymatic reactions of EcOpgD mutants were performed basically in the same way as determining the optimum pH. The concentrations of the mutants and reaction times were 0.0073–5.8 mg/mL and 0–2.5 h, respectively, depending on the mutants. Color development was performed using the MBTH method.

## Phylogenetic analysis

Consurf server^54, 55^ was used for visualizing conserved regions in the EcOpgD complex structure. Homologs that have 20%–90% amino acid sequence identities with EcOpgD were comprehensively extracted by UNIREF-90, and 150 sequences were extracted evenly from those arranged in the order of homology. Homolog sequences collected in the same way using EcOpgG as a query were used for constructing the phylogenetic tree of the MdoG superfamily. The phylogenetic tree was prepared with the maximum likelihood method using MEGA11^56^.

### Computational analysis

For analysis of each residue contribution to substrate binding, the energy effect of a mutation on substrate binding affinity was calculated as the difference between the binding free energy in the mutated and wild-type protein (ΔΔ*G*) by Discovery Studio 2018 (BIOVIA, Dassault Systèmes, San Diego, California, USA). ΔΔ*G* was defined as:

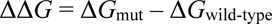

where Δ*G*_mut_ and Δ*G*_wild-type_ are the binding free energy of the mutant and wild-type enzyme, respectively. Sop_13_ was used as the substrate. Residues of EcOpgD within 5 Å of the substrate in chain B were selected for analysis. The selected residues, except alanine, were substituted to alanine, whereas alanine was substituted to glycine.

## Acknowledgments

This work was supported by Photon Factory for X-ray data collection (Proposal No. 2020G527). This work was supported in part by an academic research grant from the Mishima Kaiun Memorial Foundation. We thank Edanz (https://jp.edanz.com/ac) for editing a draft of this manuscript.

## Data availability

Atomic structure coordinates were deposited in the PDB under accession codes 8IOX, 8IP1 and 8IP2.

## Author contributions

S.M. H.N. and M.N. conceived the project and designed the experiments. S.M. expressed and purified EcOpgG and EcOpgD. S.M. and M.N. performed data collection and processed X-ray crystallographic data. K.K. performed computational analysis. H.N. provided β-1,2-glucooligosaccharides. S.M., K.K. and M.N. prepared the manuscript. H.N. and M.N. supervised the project and participated in manuscript writing. All authors contributed to the revision of the manuscript. Any correspondence should be to M.N.

