## Supplemental_Fig_Table for "Novel glycoside hydrolase family enzymes from *Escherichia coli* are associated with osmo-regulated periplasmic glucan synthesis"

**Supplemental Table 1 | Residues interacting with substrates of EcOpgD and EcOpgG**

| Subsite | Residues | |
|---|---|---|
| | EcOpgD | EcOpgG |
| −9 | (No substrate observed) | (Subsite −7) |
| −8 | (No substrate observed) | K144 (g) |
| −7 | R171, R172 | K134, E147 |
| −6 | (Subsites −4 and −7) | D146; (main) E147, Y166 |
| −5 | D173; (main) Y190 | Y56 |
| −4 | R162[a], D173[b], **R184**[c], **Y192** | Y56 (g)[d], Y83, K134[a], E147[b], **R158**, **Y166** |
| −3 | Q191, **R359** | D327, **R332** |
| −2 | N82, W441 | |
| −1 | **R184**, **D300**, **R359**, **D388***(g) | **R158**, **D273**, **R332**, **D361***(g) |
| | (main) **E385**, L387 | (main) **E358**, N360 |
| +1 | **R184**, **S195**; (main) **L178** | **R158**, **S169**; (main) **L152** |
| +2 | E445; (main) **T386** | (main) **T359** |
| +3 | **F211**, **Y182** | **F185**, **Y156** |
| +4 | **Y182**, **R197**, **E209**, **D388*** | **Y156**, **R171**, **E183**, **D361*** |
| | (main) **D388*** | (main) **D361*** |
| +5 | (Subsites +3 and +6) | (Subsites +3 and +7) |
| +6 | (Subsites +4 and +5) | E183 |
| **+7** | (No substrate observed) | (Subsites +2, +5, and +6) |

[a, b], Chemically conserved residues between EcOpgD and EcOpgG.

[c], Conserved residues between EcOpgD and EcOpgG are shown in bold letters.

[d], (g) represents hydrogen bonds with glycosidic bond oxygen atoms. Anomeric hydroxy groups are used to describe the positions of the subsites.

*Asterisks (*)* represent catalytic residues substituted with asparagine residues to obtain the substrate complexes.

*Double underlines* represent hydrophobic interactions with the substrates.

**Supplemental Table 2. Crystallographic data collection and refinement statistics of EcOpgD and EcOpgG.**

| Data set | Ligand-free EcOpgD (D378N mutant) | EcOpgD-$Sop_{13}$ (D378N mutant) | EcOpgG- $Sop_{16}$ (D361N mutant) |
|---|---|---|---|
| **Data collection** | | | |
| Beamline | KEK BL-5A | KEK BL-5A | KEK BL-5A |
| Space group | *C*222 | $P2_1$ | $C222_1$ |
| Unit cell parameters (Å) | *a* = 226.62<br>*b* = 392.36<br>*c* = 324.48 | *a* = 58.09<br>*b* = 87.10<br>*c* = 110.85<br>β = 101.13° | *a* = 62.82<br>*b* = 80.97<br>*c* = 213.23 |
| Resolution (Å)[a] | 49.14–2.95 (3.00–2.95) | 47.74–2.06 (2.11–2.06) | 48.34–1.81 (1.85–1.81) |
| Total reflections[a] | 4115005 (209213) | 434382 (21047) | 322515 (19400) |
| Unique reflections[a] | 301232 (14824) | 66585 (4130) | 50021 (2971) |
| Completeness (%)[a] | 100.0 (100.0) | 99.3 (92.6) | 99.9 (99.9) |
| Multiplicity[a] | 13.7 (14.1) | 6.5 (5.1) | 6.4 (6.5) |
| Mean $I/\sigma(I)$[a] | 13.0 (3.6) | 12.8 (2.1) | 17.2 (2.1) |
| $R_{merge}$ (%)[a] | 20.8 (87.8) | 11.6 (64.2) | 5.2 (76.2) |
| $R_{pim}$ (%)[a] | 8.4 (35.1) | 7.4 (46.6) | 3.3 (48.9) |
| $CC_{1/2}$[a] | (0.869) | (0.697) | (0.891) |
| **Refinement** | | | |
| Resolution (Å) | 49.14–2.95 | 47.74–2.06 | 48.34–1.81 |
| No. of reflections | 301231 | 63310 | 49959 |
| No. of atoms | 51848 | 9013 | 4118 |
| No. of water molecules | 666 | 566 | 122 |
| $R_{work}/R_{free}$ (%) | 19.4/23.2 | 17.5/22.4 | 21.3/25.4 |
| No. of asymmetric units | 12 | 2 | 1 |
| r.m.s.d. from ideal values | | | |
| Bond lengths (Å) | 0.0087 | 0.0084 | 0.014 |
| Bond angles (°) | 1.6025 | 1.5974 | 1.719 |
| Average *B*-factors (Å$^2$) | | | |
| Protein (chain A/B/C/…) | 34.7/40.3/37.9/35.1/<br>35.3/37.9/34.6/40.5<br>/37.9/35.2/34.7/40.5 | 24.1/24.4 | 45.1 |
| Ligand | | | |
| $Sop_n$s (chain A/B) | | 18.8/22.1 | 34.6 |
| Solvent | 32.0 | 25.8 | 34.4 |
| Ramachandran plot (%) | | | |
| Favored | 94.6 | 97.0 | 96.0 |
| Allowed | 5.2 | 3.0 | 4.0 |
| Outlier | 0.2 | 0.0 | 0 |
| **PDB entry** | 8IOX | 8IP1 | 8IP2 |

[a] Values in parentheses represent the highest resolution shell.

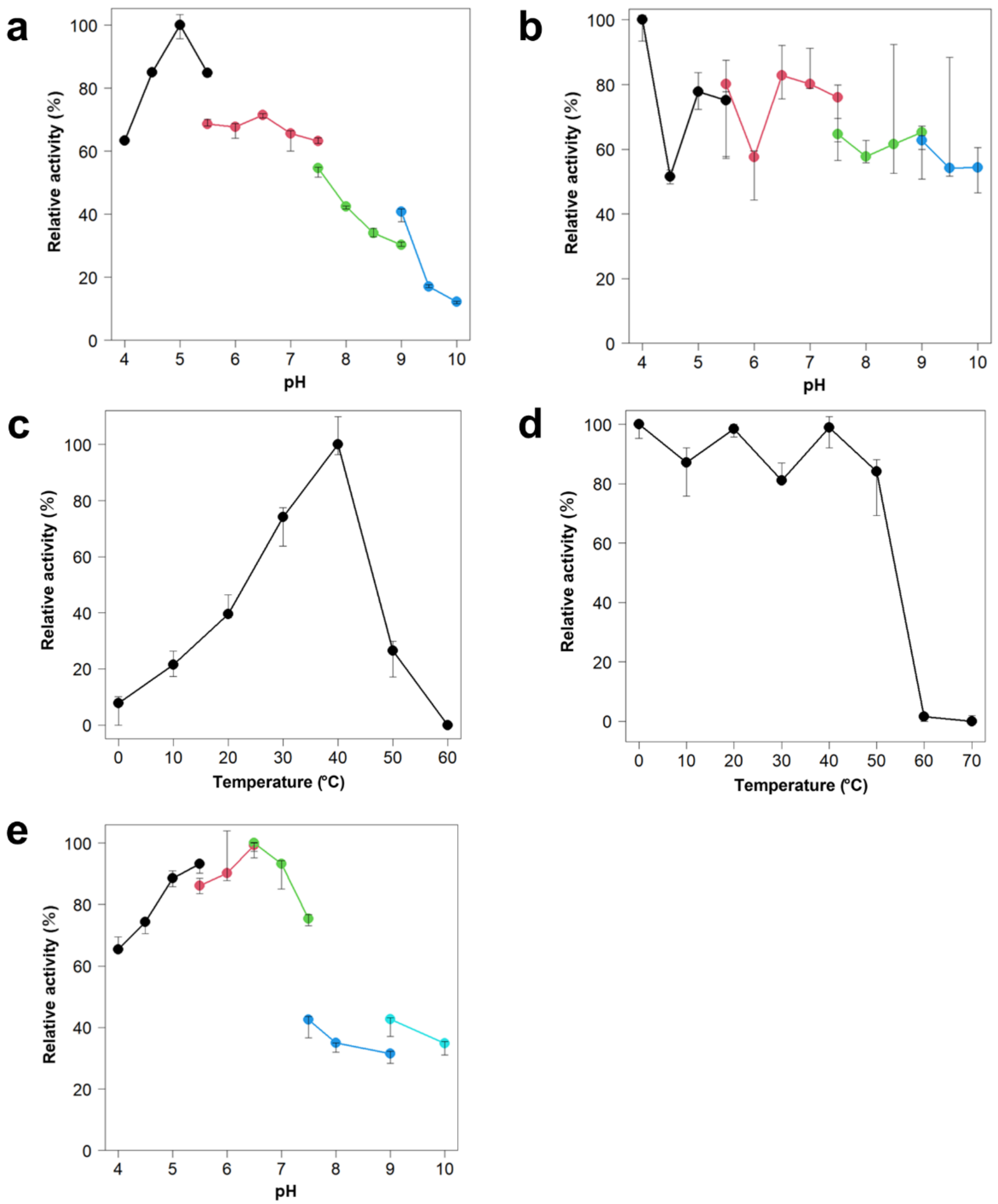


**Extended Data Fig. 1 | pH and temperature profiles.** **a**, Optimum pH of EcOpgD. Buffers used for the enzymatic reactions were sodium acetate-HCl (pH 4.0-5.5, black), Bis-Tris-HCl (pH 5.5-7.5, red), Tris-HCl (pH 7.5-9.0, green) and glycine-NaOH (pH 9.0-10, blue). **b**, pH stability of EcOpgD. Buffers used for incubating the enzyme were the same as (**a**). **c**, **d**, temperature optimum (**c**) and stability (**d**). **e**, Optimal pH of EcOpgG. Buffers used for the enzymatic reaction were sodium acetate-HCl (pH 4.0-5.5, black), MES (pH 5.5-6.5, red), MOPS (pH 6.5-7.5, green), Tris-HCl (pH 7.5-9.0, blue) and glycine-NaOH (pH 9.0-10, cyan). Medians of triplicate experiments were plotted. Other data were used to show error bars.

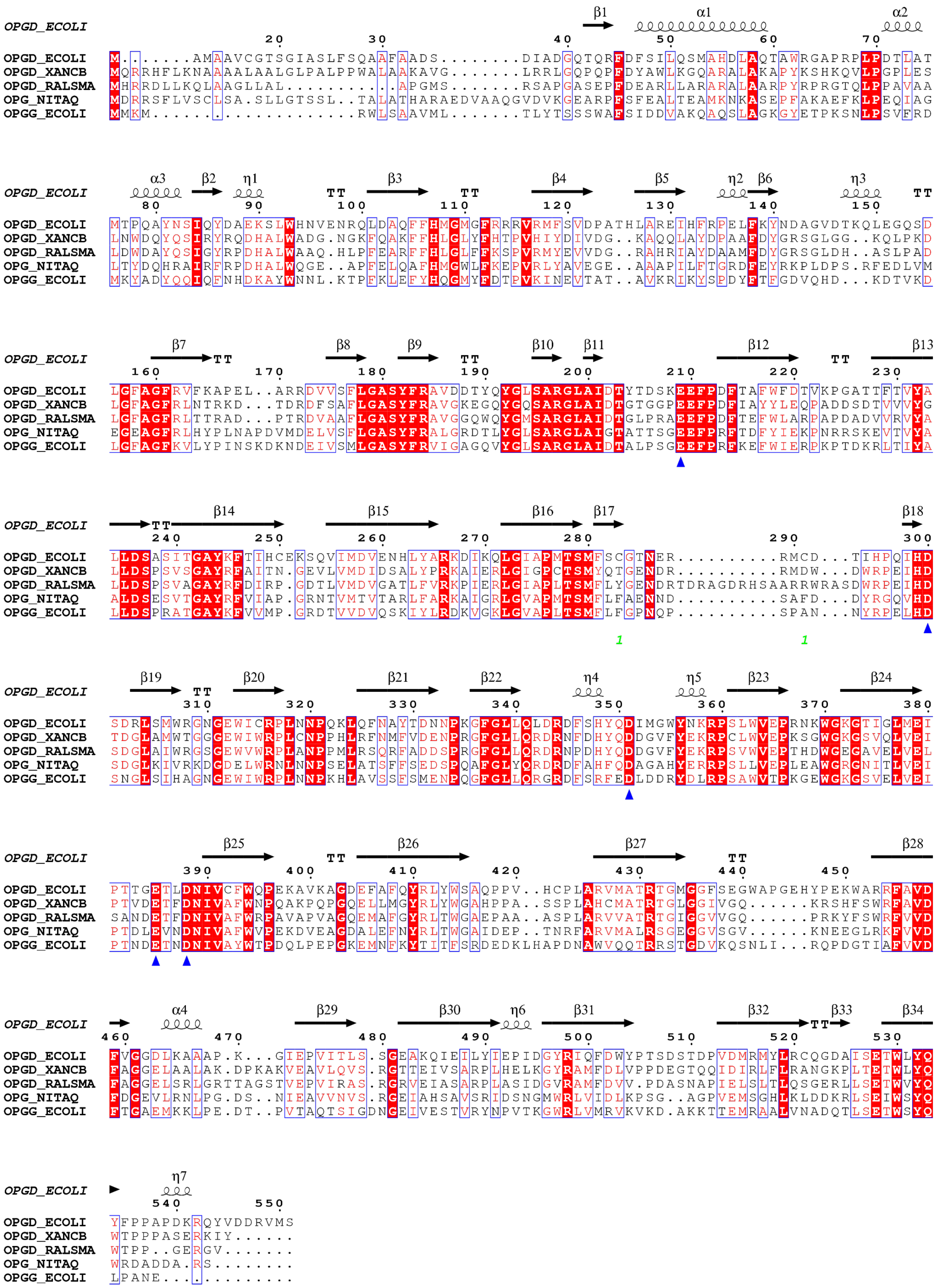


**Extended Data Fig. 2 | Multiple alignment of EcOpgD and EcOpgG.** The structure-based multiple alignment was performed by PROMALS3D[54] and visualized using the ESPript 3.0 server (http://espript.ibcp.fr/ESPript/ESPript/)[55]. Secondary structures of EcOpgD are shown above the sequences. Blue triangles are shown below all acidic residues conserved in this alignment and are located at the cleft in the ligand-free structures of EcOpgD or EcOpgG.

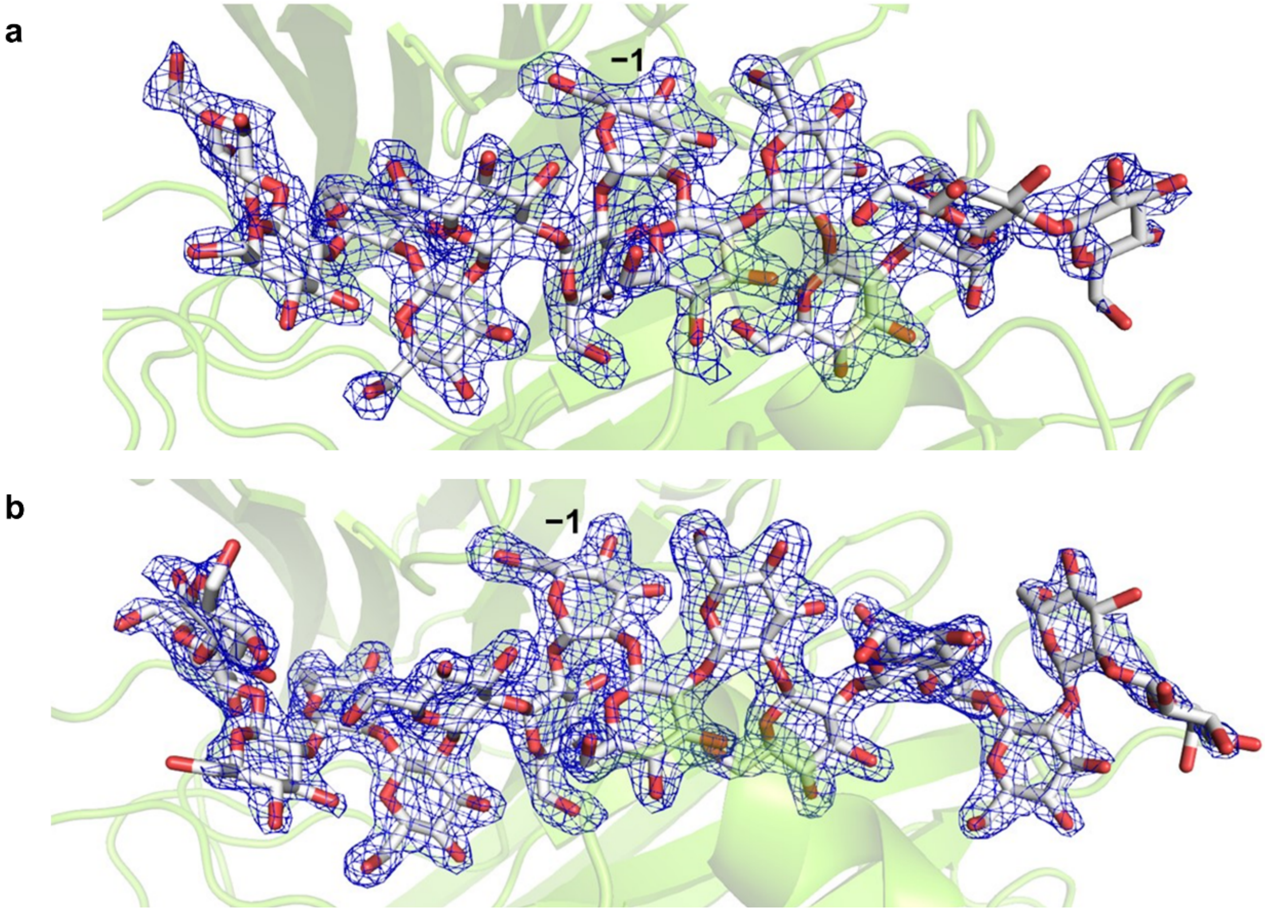


**Extended Data Fig. 3 | Electron densities of substrates in the Michaelis complex of EcOpgD and EcOpgG. a**, EcOpgD. **b**, EcOpgG. Substrates are shown as white sticks. Each chain B is shown as a semi-translucent light green. The electron densities of β-1,2-glucans are shown as $F_o$-$F_c$ omit maps by blue meshes at the 3σ contour level.

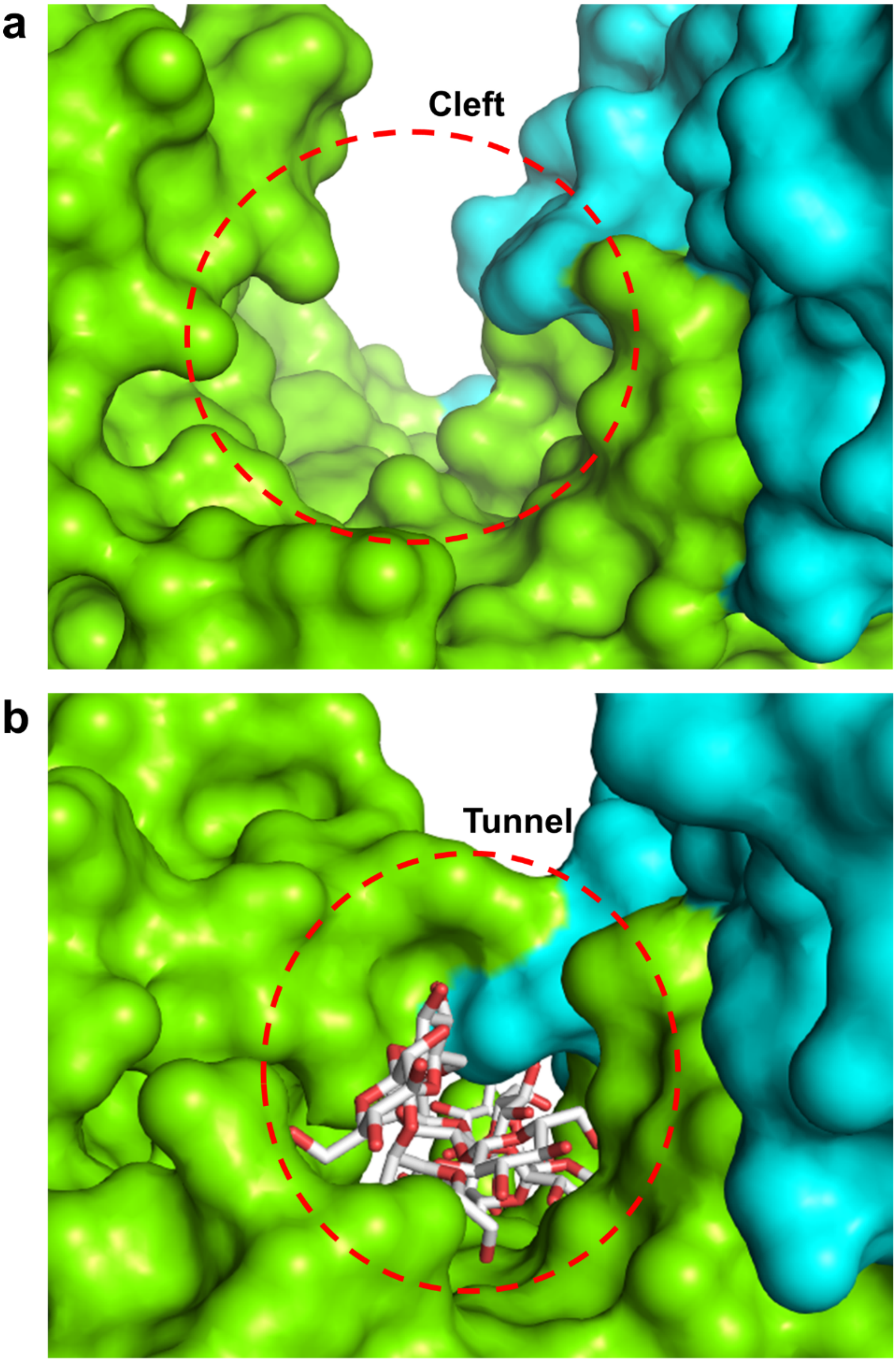


**Extended Data Fig. 4 | Substrate binding site of EcOpgD forming a cleft in the ligand-free structure and a tunnel in the Michaelis complex structure.** **a**, A cleft in the ligand-free structure. **b**, A tunnel in the Michaelis complex. Chains A and B are shown as cyan and light green surfaces, respectively. The substrate is shown as a white stick. Red dotted circles in (**a**) and (**b**) represent a cleft and a tunnel of the substrate binding sites, respectively.

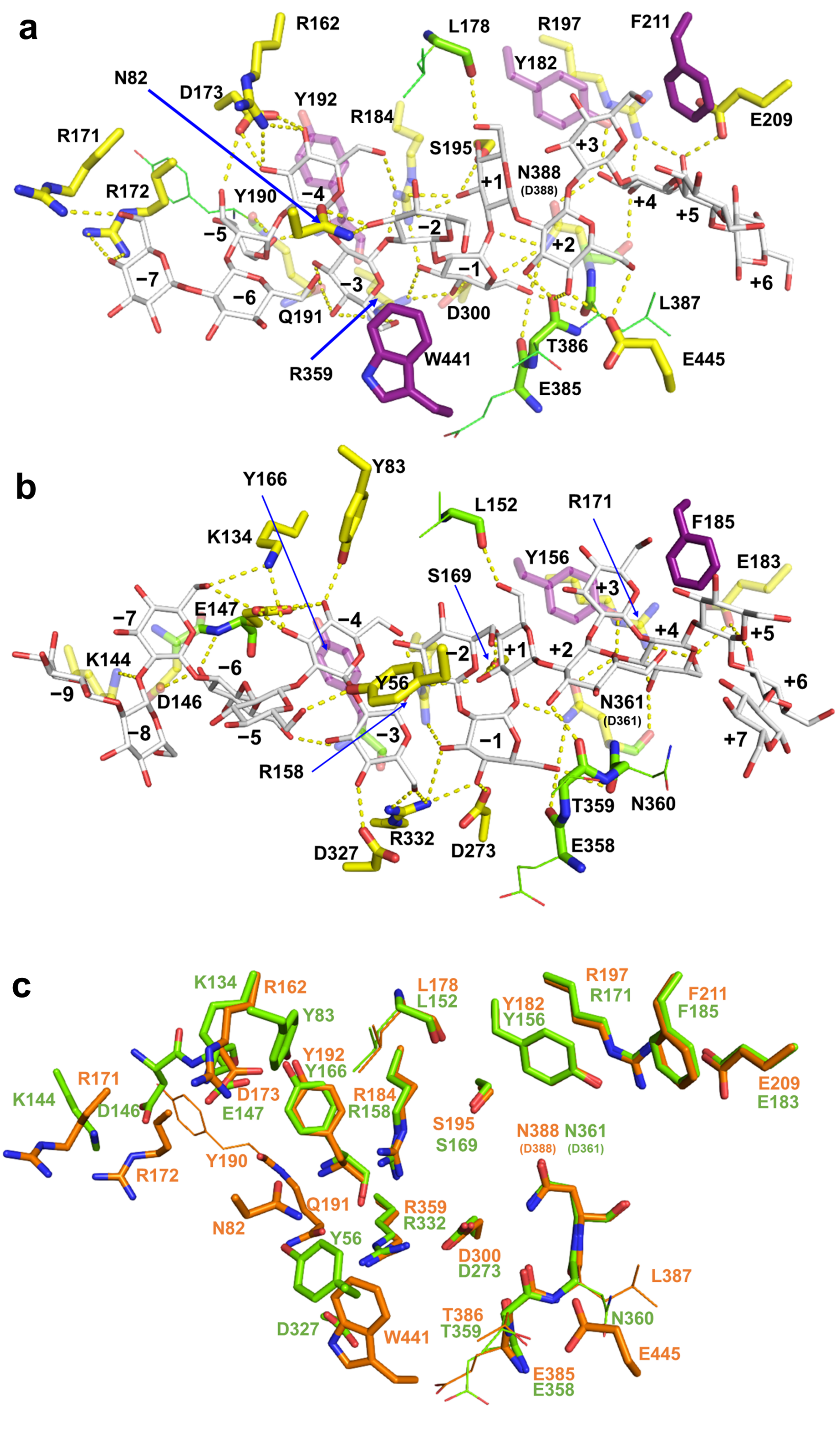


**Extended Data Fig. 5 | Substrate recognition residues in the Michaelis complexes.** **a**, **b**, Complexes of EcOpgD D388N (**a**) and EcOpgG D361N (**b**). Substrates are shown as white sticks. The main chains and side chains of residues interacting with substrates by their main chains are shown in light green sticks and lines, respectively. Residues forming hydrogen bonds with the substrates through their side chains are shown as yellow sticks. Residues forming hydrophobic interactions with the substrates are shown as purple sticks. Residues behind the substrates at this view are shown semi-translucently. Blue arrows are used to label particular amino acids. **c**, Superposition of substrate recognition residues between EcOpgD and EcOpgG. Residues of EcOpgD and EcOpgG are shown in orange and light green, respectively. The use of lines and stick models is based on (**a**, **b**).

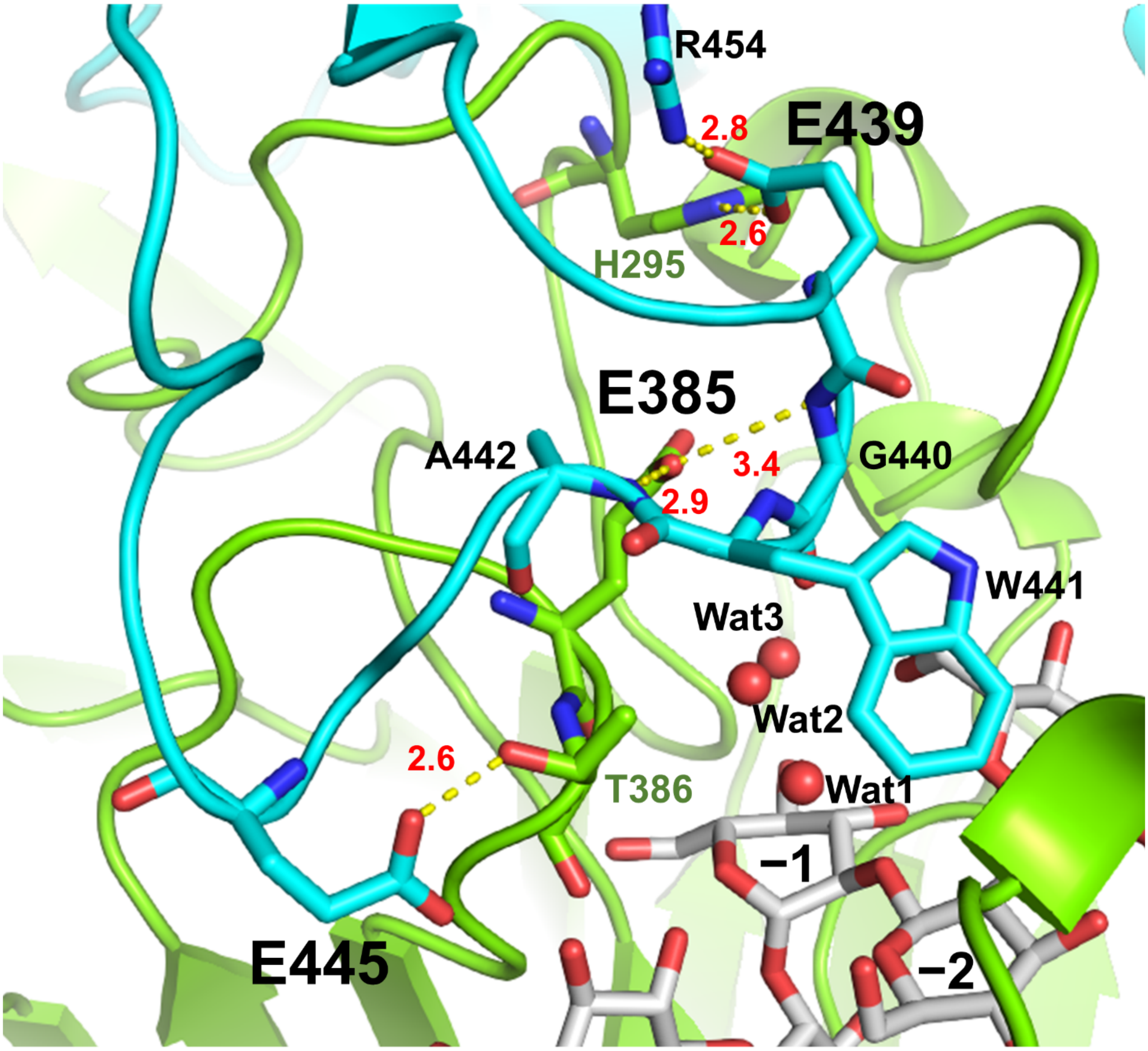


**Extended Data Fig. 6 | Interaction of Loop A with surroundings in the EcOpgD complex structure.** Chains A and B are shown in cyan and light green, respectively. Hydrogen bonds with distances (Å, red numbers) are shown as yellow dotted lines. Substrates are shown as white sticks. All residues used for the fixation of Loop A are shown as sticks. E385 interacting with the main chain nitrogen atoms of G440 and A442, E439 interacting with H295 and R454, and E445 interacting with T386 are labeled with large bold letters. Residues in chain B are labeled in green letters. Wat1, Wat2 and Wat3 are shown as red spheres.

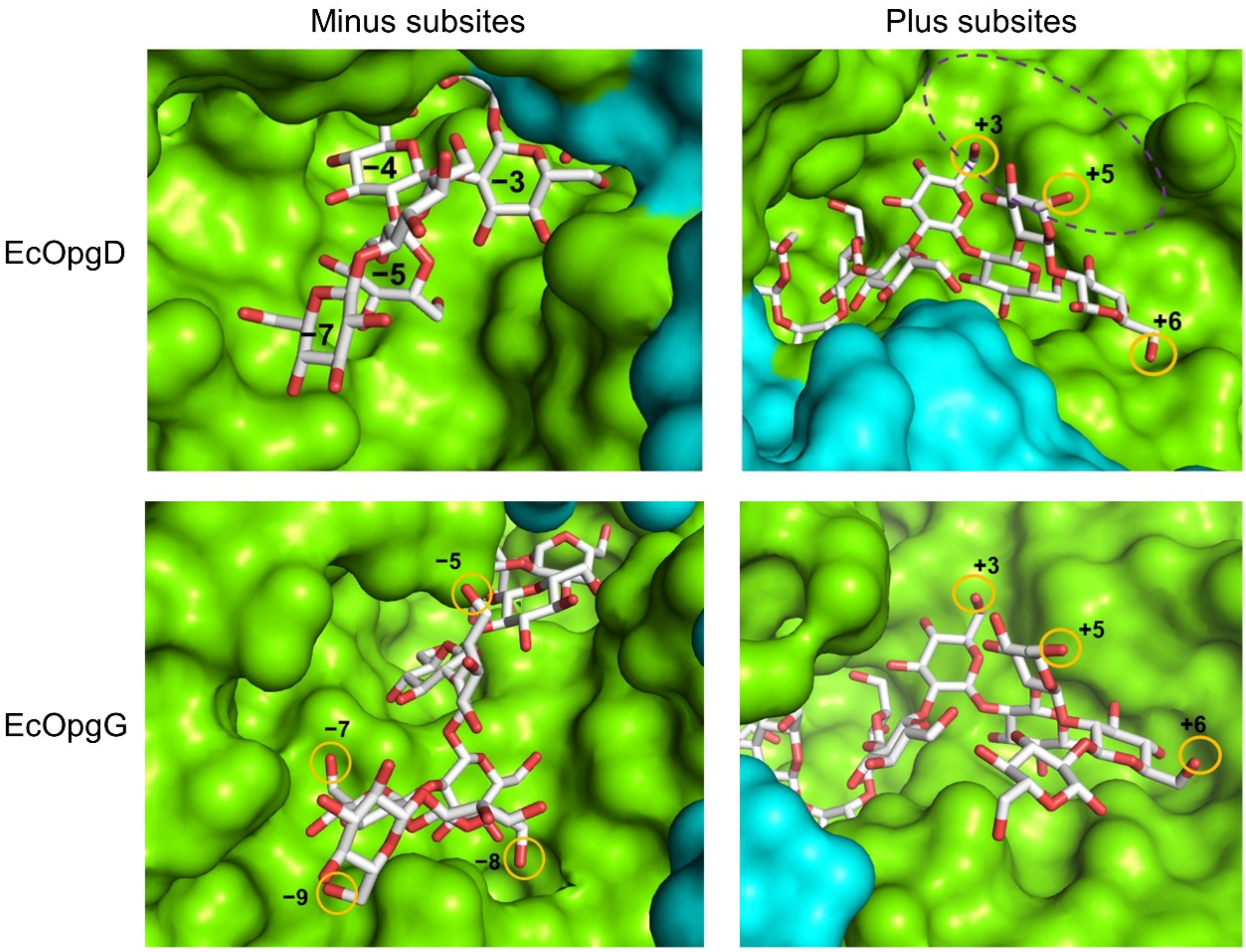


**Extended Data Fig. 7 | Spaces for β-1,6-side chains at minus and plus subsites of EcOpgD and EcOpgG.** Chains A and B are shown in cyan and light green surfaces, respectively. Substrates are shown as white sticks. The 6-hydroxy groups of Glc moieties at the subsites, which clearly have sufficient space for Glc side chains, are indicated by orange circles. The large space of EcOpgD that accommodates β-1,6-linked side chains is indicated as a dotted circle.

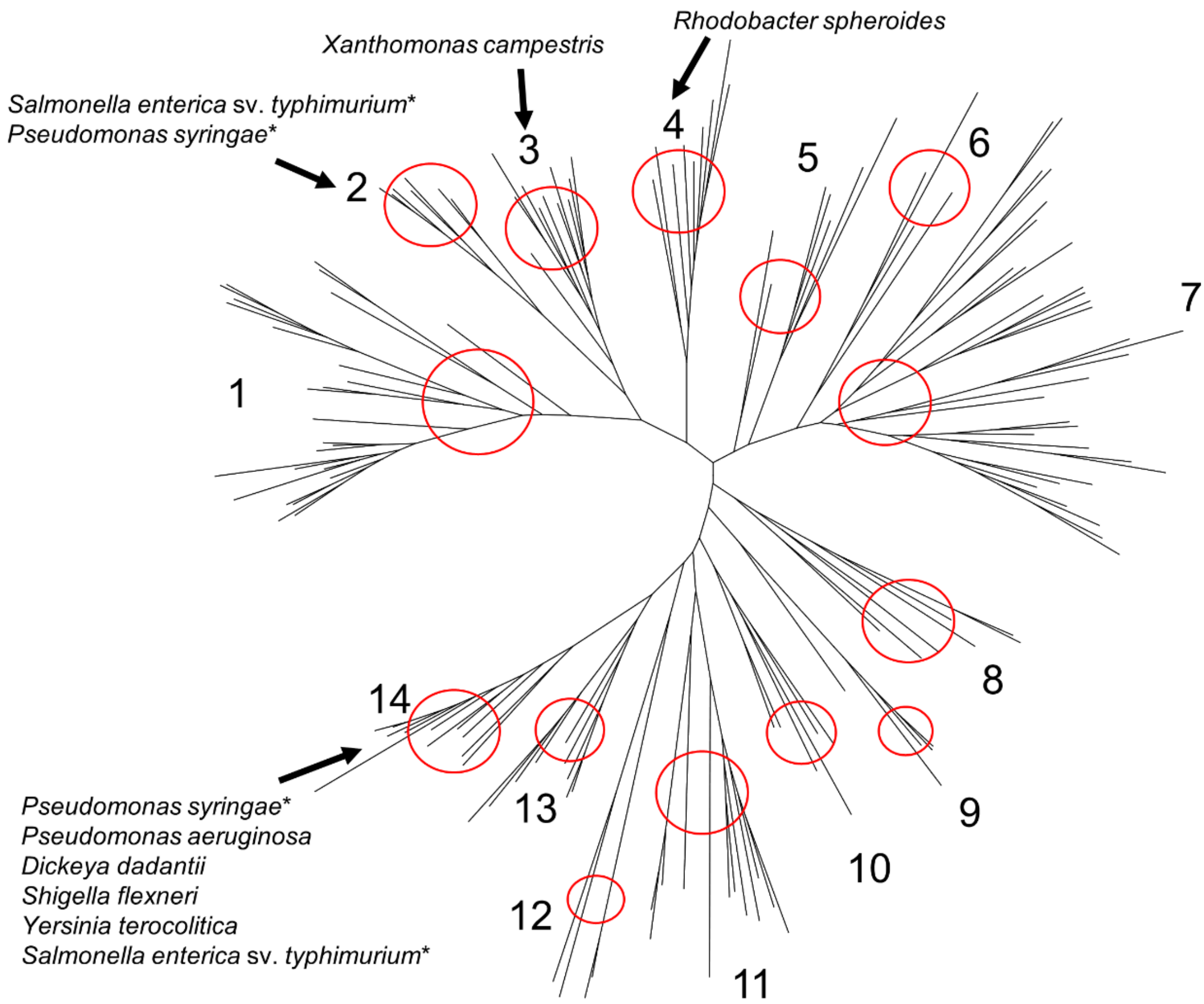


**Extended Data Fig. 8 | Phylogenetic tree of the MdoG superfamily.** Each clade is indicated by a red circle with a clade number. Arrows indicate clades, including homologs from Gram-negative bacteria whose phenotypes of OPG-related genes knockout mutants have been investigated. Asterisks represent species possessing two homologs.

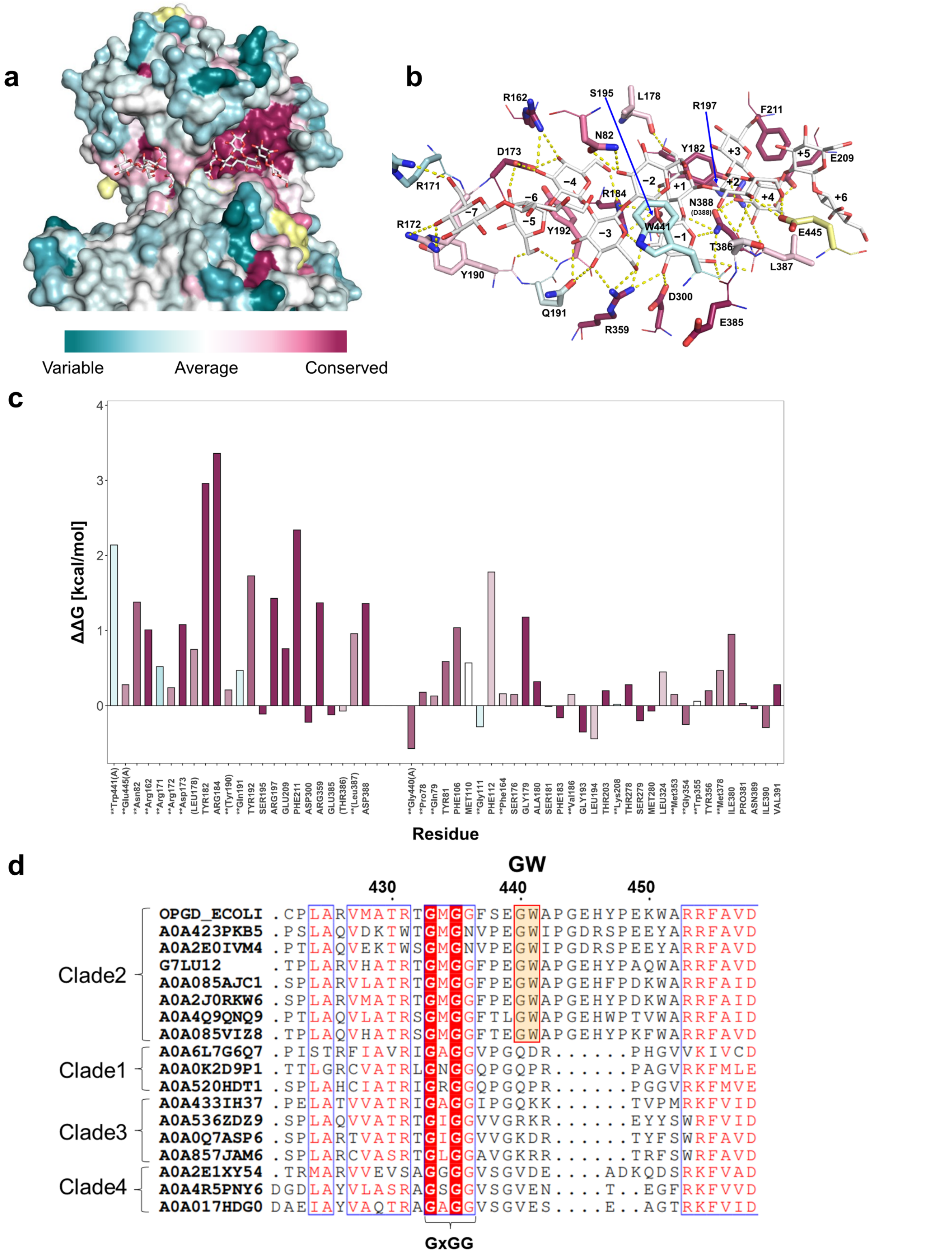


**Extended Data Fig. 9 | Conservation of residues among the MdoG superfamily.** a, b, Conserved residues of EcOpgD. Surface model (a) and substrate recognition residues (b) of the EcOpgD-β-1,2-glucan complex colored based on the color bar corresponding to conservation scores. The conservation scores were calculated by Consurf using EcOpgD. c, Contribution of residues for substrate binding. Residues shown on the left and right sides are substrate recognition residues and residues in close proximity to the substrate rather than substrate recognition, respectively. The colors used for the bars are the same as (a, b). Parentheses represent the main chains that recognize the substrate. Residues in chain A are indicated with (A). Asterisks denote residues that are not conserved in EcOpgG. d, Multiple alignment of clades 1-4. Multiple alignment of homologs in clades 1-4 was visualized using the ESPript 3.0 server (http://espript. ibcp.fr/ESPript/ESPript/)55. The homologs are represented by UniProt accession numbers. Residue numbers of EcOpgD are shown above the alignment.
